# Functional diversity of the *TP53* mutome revealed by saturating CRISPR mutagenesis

**DOI:** 10.1101/2023.03.10.531074

**Authors:** Julianne Funk, Maria Klimovich, Evangelos Pavlakis, Michelle Neumann, Daniel Drangenstein, Maxim Noeparast, Pascal Hunold, Anna Borowek, Dimitrios-Ilias Balourdas, Katharina Kochhan, Nastasja Merle, Imke Bullwinkel, Michael Wanzel, Sabrina Elmshäuser, Andrea Nist, Tara Procida, Marek Bartkuhn, Katharina Humpert, Marco Mernberger, Rajkumar Savai, Andreas C. Joerger, Thorsten Stiewe

## Abstract

The tumor suppressor gene *TP53* is the most frequently mutated gene in various cancers. Unlike other tumor suppressors, *TP53* is mostly hit by missense mutations, of which more than 2,000 have been described in cancer patients. To take advantage of *TP53* mutation status for personalized therapy, a deeper knowledge of the functional ramifications of specific mutations is required as evidence of the functional heterogeneity of mutant p53 proteins mounts. Here, we report on a CRISPR-based saturation mutagenesis screen of 9,225 variants expressed from the endogenous *TP53* gene locus of a cancer cell. By tracking changes in the abundance of individual variants in response to specific p53-pathway stimulation, we were able to construct high-resolution functional activity maps of the *TP53* mutome, covering ∼94.5% of all cancer-associated missense mutations. The results demonstrate the impact of individual mutations on tumor cell fitness with unprecedented precision and coverage, even revealing underlying mechanisms such as apoptosis. The high discriminatory power also resolves subtle loss-of-function phenotypes and highlights a subset of mutants as particularly promising targets for pharmacological reactivation. Moreover, the data offer intriguing insight into the role of aberrant splicing and nonsense-mediated mRNA decay in clearing truncated proteins due to not only nonsense, frameshift, and splice-site mutations but also missense and synonymous mutations. Surprisingly, no missense mutation provided an immediate proliferative advantage over a null mutation. Nonetheless, cells with a missense, but not null mutations, acquired pro-metastatic properties after prolonged growth in mice, emphasizing the significance of mutant p53-directed clonal evolution in the progression of tumors towards metastasis.

## Introduction

p53, a master regulatory transcription factor, reduces the proliferative fitness of cancer cells, thereby limiting their clonal expansion. This is achieved through a constantly expanding array of mechanisms, including cell-cycle arrest, senescence, and apoptosis^1^. Mutations in the p53-encoding gene *TP53* are observed in about half of all cancer patients and, as germline mutations, lead to the hereditary Li-Fraumeni cancer susceptibility syndrome^2^. In many cancers, *TP53* mutations correlate with enhanced metastasis and aggressiveness, decreased response to treatment, and thus poor prognosis^3^. However, despite the high frequency of *TP53* mutations in cancer patients and their prognostic implications, the use of *TP53* mutation status in clinical decision-making is limited because of the complexity of the *TP53* mutome, i.e., the broad spectrum of functionally diverse mutations, which makes it difficult to predict the pathogenicity and clinical consequences of an individual mutation.

The mutome of a cancer gene, and in particular that of *TP53*, arises from biased mutational processes and further editing by evolutionary pressures that deplete immunogenic neoantigens and select for mutations that promote tumor cell fitness^4-8^. Unlike other tumor suppressor genes, *TP53* mutations are, in the majority of cases, missense mutations that result in the accumulation of full-length p53 proteins with a single amino acid alteration. To date, more than 2,000 different missense mutations have been described in cancer samples^9^. No single mutation occurs at a frequency greater than 6%, and the ten most common (and also most studied) ‘hotspot’ mutants jointly account for only ∼30%. In turn, the vast majority (∼70%) of patients with *TP53* mutations have one of >2,000 different, poorly characterized non-hotspot mutations^9^.

We can distinguish three principal consequences of a *TP53* missense mutation: loss of wild-type function (LOF), dominant-negative effects (DNE), and gain-of-function (GOF) properties^10,11^. LOF refers to a variable, more or less extensive reduction in tumor suppressive functions. The DNE refers to inhibition of wild-type p53 by mutant p53 (mutp53), while GOF refers to neomorphic properties conveyed by amino acid sequence alterations^12-18^. GOF can promote tumor metastasis, confer drug resistance and immune escape, and generate an oncogene-like addiction that makes tumor cells dependent on the presence of the mutant protein. GOF appears to be highly context-specific, dependent on protein level and the precise amino acid change^12-16^.

Given the high prevalence of *TP53* missense mutations in cancer patients and the complexity of the mutome, an in-depth understanding of single mutants and their functional consequences is crucial to better utilize p53 status clinically for personalized treatment decisions. However, most p53 mutations are rare, making it statistically challenging to infer their clinical implications from patient cohorts. High-throughput screens in isogenic cellular models are therefore a valuable tool for functionally annotating the *TP53* mutome. The first comprehensive and systematic analysis was based on a cDNA library of 2,314 missense variants screened for transcriptional activity in a yeast-based reporter system with p53-responsive elements of 8 selected p53 target genes^7^. While this study revealed a widespread loss of transcriptional activity for the hotspot and many non-hotspot cancer mutants, it also highlighted an unexpected heterogeneity with many non-hotspot mutants retaining considerable residual transactivation potential at one or more response elements^7^. Due to the absence of p53, yeast cells lack the p53 control network, which includes key p53 regulators such as Mdm2, as well as other enzymes that fine-tune p53 function through post-translational modifications. In subsequent screens, mutp53 cDNA libraries were therefore analyzed in human cells^4,6,8^. However, cDNA-based expression screens have inherent limitations, among them non-physiological expression driven by heterologous promoters, the absence of post-transcriptional control through miRNA-UTR interactions, and the lack of (alternative) splicing. Moreover, as these studies focused on the consequences of introducing p53 variants into p53-deficient cell lines, they did not systematically assess the clinically most relevant impact of p53 mutations on tumor cell responses to cancer treatments, such as radiation, chemotherapy or targeted therapies.

Here, we have leveraged the power of CRISPR/Cas9-mediated gene editing through precise homology-directed repair to introduce a panel of 9,225 variants, comprising approximately 94.5% of all *TP53* cancer mutations, into cancer cells with a wild-type *TP53* gene locus and a prototypical p53 response. Unlike previous cDNA overexpression screens, CRISPR-based *TP53* gene editing preserves the physiological regulation of expression through its endogenous regulators, including promoters, enhancers, splice factors, and miRNA binding sites in the UTRs. We have measured the impact of all variants on proliferative fitness following highly specific p53 pathway activation with Mdm2 inhibitors and confirmed similar but quantitatively less pronounced fitness effects for other p53-activating stimuli, including radiation, chemotherapy and starvation. The fitness effects of mutants correlated with their abundance in cancer patients, evolutionary conservation, and known structure-function relationships. In contrast to cDNA overexpression screens, CRISPR-mediated expression from the endogenous *TP53* gene locus allowed for accurate annotation of partial loss-of-function and splice mutations and has demonstrated widespread elimination of frameshift, nonsense and splice mutations by nonsense-mediated mRNA decay. Notably, none of the missense variants resulted in a fitness gain beyond that of null-mutations. Surprisingly, cells with missense but not nonsense mutations stabilized the mutant protein and developed metastatic properties when grown in mice, indicating that GOF effects are not an intrinsic property of the *TP53* mutations but rather a result of mutant p53-directed tumor evolution.

## Results

### Isogenic model for *TP53* mutagenesis by CRISPR-HDR

In order to compare the functional impact of *TP53* variants in an isogenic setting, we chose HCT116 colorectal carcinoma cells as our model system. These cells are wild-type for *TP53* and show a prototypical p53 response, making them a well-established model for investigating the mechanistic details of p53-mediated tumor suppression. Additionally, HCT116 cells have a homozygous *MLH1* nonsense mutation, resulting in a mismatch repair deficiency, which facilitates gene targeting through homologous recombination^19,20^. To ensure unambiguous genotype-phenotype correlations, we haploidized the cells for *TP53* by deleting intronic splicing branch points in one of the two *TP53* alleles (Δ allele, Fig. 1a and Supplementary Fig. 1a-c). To prevent potential bias from the anti-proliferative p53 response triggered by CRISPR/Cas9 nucleases during gene editing^21-23^, we silenced expression from the remaining copy reversibly by inserting a LoxP-flanked transcriptional stop cassette (Lox-Stop-Lox, LSL) into intron 4 (HCT116 LSL/Δ). The LSL cassette included an EGFP expression cassette for monitoring and a mutated puromycin N-acetyltransferase (*pac*) gene that could be repaired during editing for later selection. The LSL allele was specifically cleaved by CRISPR/Cas9 nucleases targeting the intronic regions deleted on the Δ allele. Specific mutations of interest were introduced into the cut LSL allele by homology-directed repair (HDR) with co-transfected donor vectors harboring the mutations, a mutated protospacer adjacent motif (PAM) to prevent re-cutting, and the LSL-cassette with a repaired *pac* gene. Correctly HDR-edited cells (HCT116 LSL-mut/Δ) were selected with puromycin and infected with Cre recombinase expressing adenovirus (AV-Cre) to induce monoallelic p53 variant expression (HCT116 mut/Δ).

**Fig. 1:**
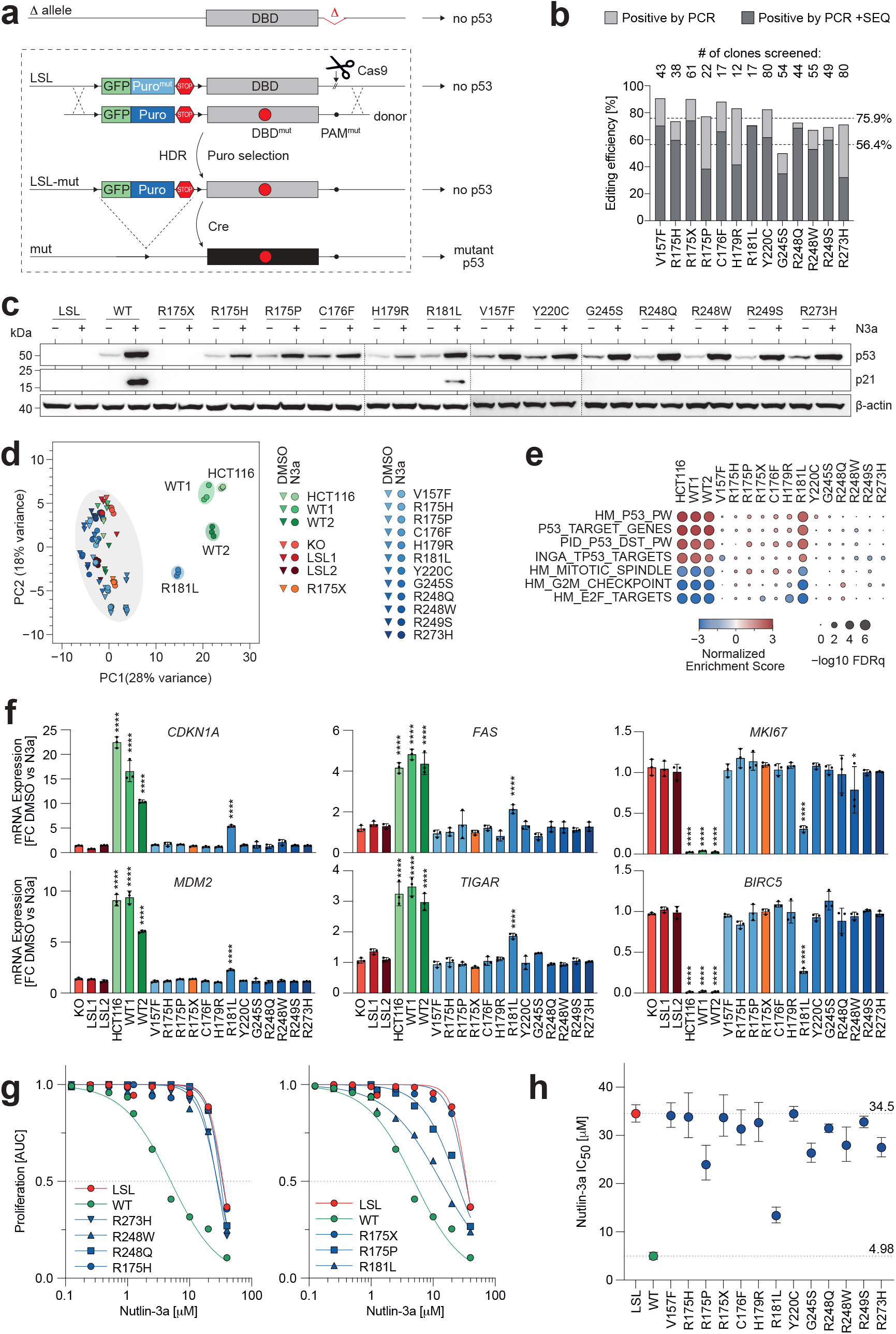
Panel of single *TP53* mutations in HCT116 cell lines. **a**, Scheme for CRISPR/Cas9-mediated *TP53* mutagenesis via homology-directed recombination (HDR) in HCT116 LSL/Δ cell line. LSL, loxP-stop-loxP; DBD, DNA-binding domain. **b**, Editing efficiency as percentage of single-cell clones that contain a targeted integration of the donor and the desired mutation analyzed by PCR and sequencing, respectively. Shown are results for single mutations and the mean across the panel. **c**, Western blot demonstrating mutant p53 protein expression in HCT116 clones after Cre-mediated excision of the LSL cassette in absence and presence of 10 µm N3a. **d**, Principle component analysis based on RNA-seq data of indicated cells clones ±N3a. **e**, Gene set enrichment analysis for p53-related gene expression signatures comparing indicated N3a and DMSO-treated cell clones. **f**, mRNA-expression changes of representative p53-activated and repressed genes following N3a-treatment in cell clones. Shown is the mean ±SD. Data points indicate data from replicate RNA-seq datasets (n=3). ****, p<0.0001; *, p<0.05; two-way ANOVA with Sidak’s post-hoc multiple comparisons test. **g-h**, Proliferation of *TP53*-mutant cell clones in presence of increasing concentrations of N3a analyzed by real-time live cell imaging. **g**, Area under the proliferation curve relative to untreated. **h**, 50% inhibitory concentration (mean IC_50_ and 95% CI, n=3) for N3a with p53-null (LSL, red) and wild type (WT, green) as reference.

We validated the performance of *TP53* editing in HCT116 LSL/Δ cells by introducing a panel of *TP53* variants, including some of the most frequent cancer mutations (V157F, R175H, C176F, H179R, Y220C, G245S, R248Q, R248W, R249S, R273H) with known LOF, the partial LOF (pLOF) mutations R175P and R181L, the nonsense mutation R175X and the wild-type (WT) for reference. Single-cell clones were analyzed and showed correct editing in 75.9% by PCR and 56.4% by DNA sequencing (Fig. 1b). Excision of the LSL cassette with AV-Cre resulted in comparable protein expression levels of WT and all p53 missense mutants, which was further boosted by inhibition of Mdm2 with Nutlin-3a (N3a) (Fig. 1c and Supplementary Fig. 1d). Consistent with the expected LOF of many cancer mutants, N3a induced expression of p21/CDKN1A protein and characteristic p53 gene expression signatures only in WT and R181L mutant cells (Fig. 1c-f). Real-time live cell imaging of N3a-treated cells demonstrated a strong reduction in proliferative fitness in WT cells, which was diminished by pLOF mutations and completely abrogated by LOF missense and nonsense mutants (Fig. 1g,h and Supplementary Fig. 1e). Notably, different R175H single-cell clones showed only little variation in p53 expression, transcriptional activity, and N3a response (Supplementary Fig. 1f-h). These data validate HCT116 LSL/Δ cells as an efficiently editable cellular model, suitable for exploring the functional impact of *TP53* variants in a high-throughput manner.

### Saturating R175 screen reveals functional differences between missense variants

Leveraging the editability of HCT116 LSL/Δ cells, we conducted a saturating mutagenesis of codon R175, the most frequently mutated codon in cancer, represented by R175H. We generated a donor library consisting of 27 plasmids, each containing a distinct variant of amino acid substitution (missense, mis), in-frame (if) and frameshift (fs) deletions (del), and insertions (ins), as well as a nonsense mutation (non) and several silent/synonymous (syn) mutations. LSL/Δ cells were co-transfected with a *TP53* intron 5 targeting CRISPR nuclease and the donor library, and edited LSL-mut/Δ cells were selected with puromycin, maintaining an average coverage of at least 1,000 independently edited cells per variant (Fig. 2a).

**Fig. 2:**
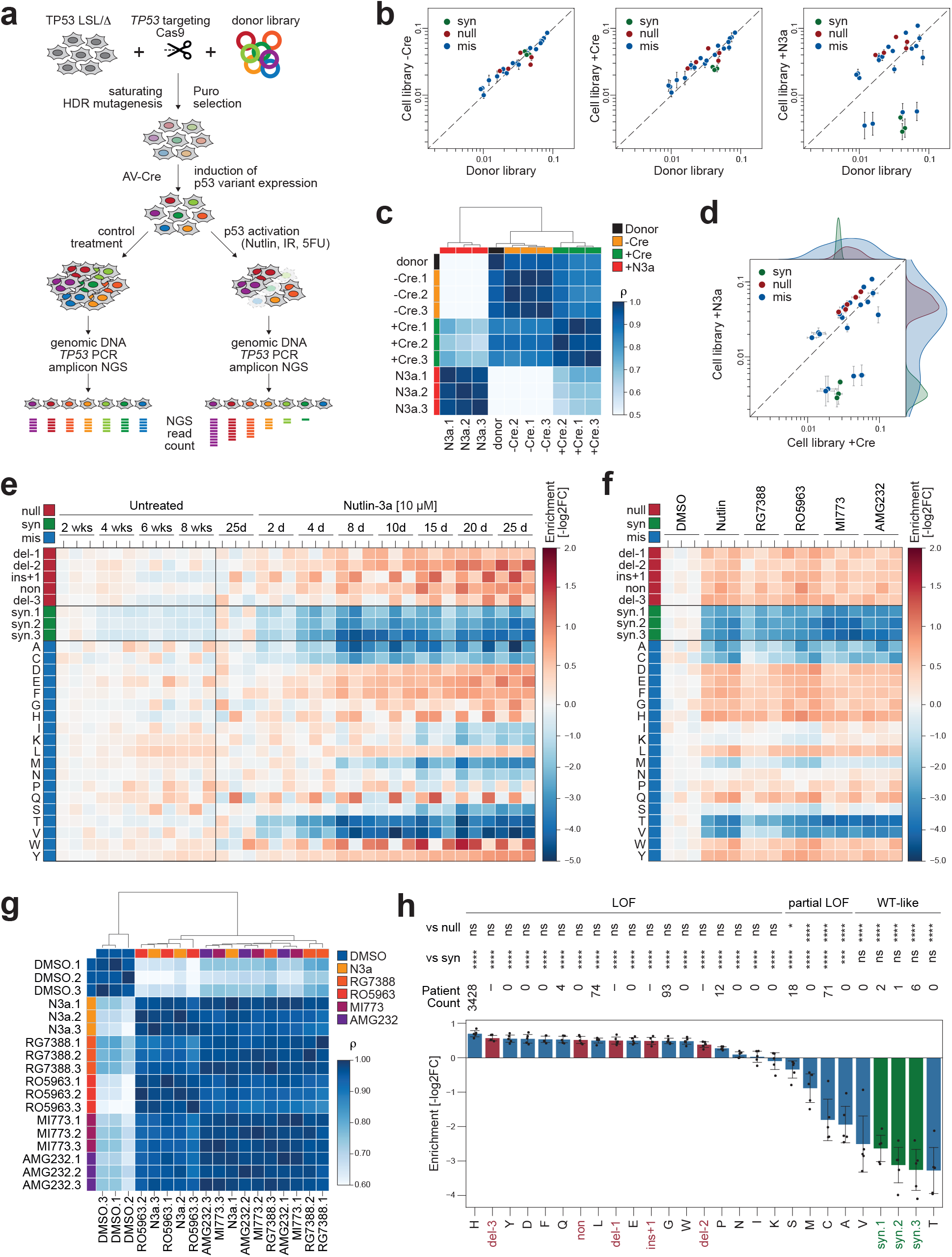
Saturating mutagenesis of *TP53* codon R175. **a**, Scheme for CRISPR/Cas9-mediated saturating mutagenesis via homology-directed recombination (HDR) in HCT116 LSL/Δ cell line and analysis of p53-mediated stress responses by next generation sequencing (NGS). Puro, puromycin; IR, ionizing radiation; 5-FU, fluorouracil. **b** and **d**, Quality control plots illustrating correlation of variant abundance between donor (plasmid) library and variant cell libraries before and after Cre recombination (-Cre and +Cre) and following 8 days of N3a treatment (+N3a). Shown is the mean ±SD abundance (n=3 biological replicates) for synonymous (syn, green), null (red) and missense (mis, blue) variants. Kernel density estimation plots illustrate separation of variants following N3a treatment. Dashed line, line of identity. **c**, Heatmap showing pair-wise correlation coefficients (ρ, Spearman). Dendrogram shows hierarchical clustering of samples using average linkage and Euclidean distance. **e**, Heatmap showing the temporal changes of variant abundance in absence or presence of N3a (n=3 biological replicates per condition). Enrichment or depletion is shown as the -log2 fold change versus the mean of the early untreated replicates. **f** and **g**, Response to Mdm2/Mdmx inhibitors. **f**, Heatmap of variant enrichment/depletion after 8 days of treatment (n=3 biological replicates per condition). **g**, Heatmap showing pair-wise correlation coefficients (ρ, Spearman) with hierarchical clustering of samples using average linkage and Euclidean distance. **h**, Bar plot of variant enrichment/depletion shown as mean±SD of the five tested Mdm2/Mdmx inhibitors. Each data points represents the enrichment for one compound (median of 3 replicates). Null mutations are highlighted in red, synonymous variants in green. Statistical significance was tested by one-way ANOVA. Reported are p-values from Dunnett’s post-how multiple comparisons test for each variant versus the mean of all synonymous or all null variants. For each variant, the patient sample count in the UMD mutation database is stated.

The editing performance was validated through targeted amplicon sequencing of the R175-containing exon 5. The results showed a strong correlation between the variant distributions in the donor plasmid and cell libraries across independent biological replicates (transfections) (Fig. 2b, c and Supplementary Table 1). This correlation was maintained even after activation of p53 variant expression by AV-Cre (Fig. 2b,c), indicating high reproducibility and absence of significant variant bias during the editing or AV infection process. Upon treatment with N3a, the variant distribution changed significantly and lost correlation with the untreated cell libraries (Fig. 2b-d).

Syn variants became strongly depleted and clearly separated from non/fs variants (grouped as ‘null’ mutations), while mis variants showed a highly variable mutation-specific response (Fig. 2d,e). To quantify the relative enrichment or depletion of variants within the cell library upon treatment, we calculated an enrichment score (ES), defined as the negative log2-transformed fold change compared to the control treatment (Fig. 2e). In the absence of N3a, the variant distribution in the Cre-recombined cell libraries remained stable for at least eight weeks, displaying only minor depletion of syn variants (Fig. 2e). The N3a response increased with time and N3a concentration and displayed a similar pattern across a range of different Mdm2 and Mdmx inhibitors (Fig. 2e-g and Supplementary Fig. 2a), validating that the measured fitness effects reflect the consequences of p53 activation rather than p53-independent (off-target) effects of N3a. When comparing all mis variants (Fig. 2h), we observed three classes: ‘LOF’ variants like R175H with an ES significantly different from WT and syn mutations, ‘pLOF’ variants with an ES significantly different from both syn and null mutations, and ‘WT-like’ variants that were statistically indifferent from syn mutations. All recurrent R175 mutations in cancer patients fall into the LOF and pLOF class, and the most frequent ones are uniformly classified as LOF, demonstrating the power of the CRISPR mutagenesis screen to correctly identify pathogenic mutations.

Over the years, several compounds have been described that may restore tumor suppressive activity to p53 mutants^3,24,25^. However, the majority of these compounds display considerable p53-independent toxicity, which contributes to, if not determines, their therapeutic effects^24^. In order to test the capability of two distinct compounds, the alkylating agent APR-246 and the metallochaperone ZMC1^26,27^, to reactivate mutant p53 in an isogenic setting, we utilized the R175-mutant HCT116 cell pools. Drug-induced reactivation of tumor suppressor activity is expected to reduce the proliferative fitness of cells and selectively deplete cells with reactivatable missense mutants. However, upon treatment with APR-246 or ZMC1, we observed no changes in the abundance of any mutation, including the top target R175H (Supplementary Fig. 2b-e). Even when treating the cell pools with a combination of APR-246 or ZMC1 and the Mdm2 inhibitor N3a, the abundance changes were indistinguishable from treatment with N3a alone. This absence of specific depletion of R175H or other missense mutants strongly indicates that APR-246 and ZMC1 are not capable of reactivating R175 missense mutants sufficiently to reduce proliferative fitness. This result is consistent with studies suggesting that the anti-tumor activity of these compounds may rely more on interference with redox homeostasis than reactivation of mutant p53^28-31^.

To examine the cell type-specificity of the results, we performed a similar experiment in non-small cell lung cancer cell line H460, which carries 3 wild-type copies of *TP53*. Analogously to HCT116 LSL/Δ cells, we made the expression of one wild-type allele conditional and deleted the remaining two copies, resulting in H460 LSL/Δ/Δ cells (Supplementary Fig. 3a,b). After confirming the effectiveness of the editing process (Supplementary Fig. 3c,d), we introduced the R175 variant library into the editable LSL allele, activated expression with Cre, and measured the response of the H460 mut/Δ/Δ cell library to N3a treatment as described for HCT116 cells (Supplementary Fig. 3e,f). The N3a-induced changes in the abundance of individual p53 variants were significantly correlated between H460 cells and HCT116 cells (ρ=0.969, p<0.0001), indicating that the fitness effect of mutations is highly conserved across different cell types.

We further evaluated the relationship between the response of the R175 variant to specific p53 pathway activation with N3a and the response to other key stress signals in the p53 pathway, such as DNA damage and nutrient deprivation. To this end, we treated the R175 cell library with varying doses of X-irradiation (IR) or 5-fluorouracil (5-FU), starved them in Hank’s Balanced Salt Solution (HBSS) or deprived them selectively of glucose or glutamine (Fig. 3a). Under all conditions, the fitness effects correlated significantly with the pattern observed upon N3a treatment. However, the overall effects were less pronounced (Fig. 3b), likely due to the contribution of p53-independent mechanisms to the analyzed stress responses, which attenuates the impact of p53 variants. These results indicate that Mdm2 inhibitors, because of their selectivity for the p53 pathway, are more effective than other p53-activating stimuli in discriminating the functional differences of p53 variants.

**Fig. 3.**
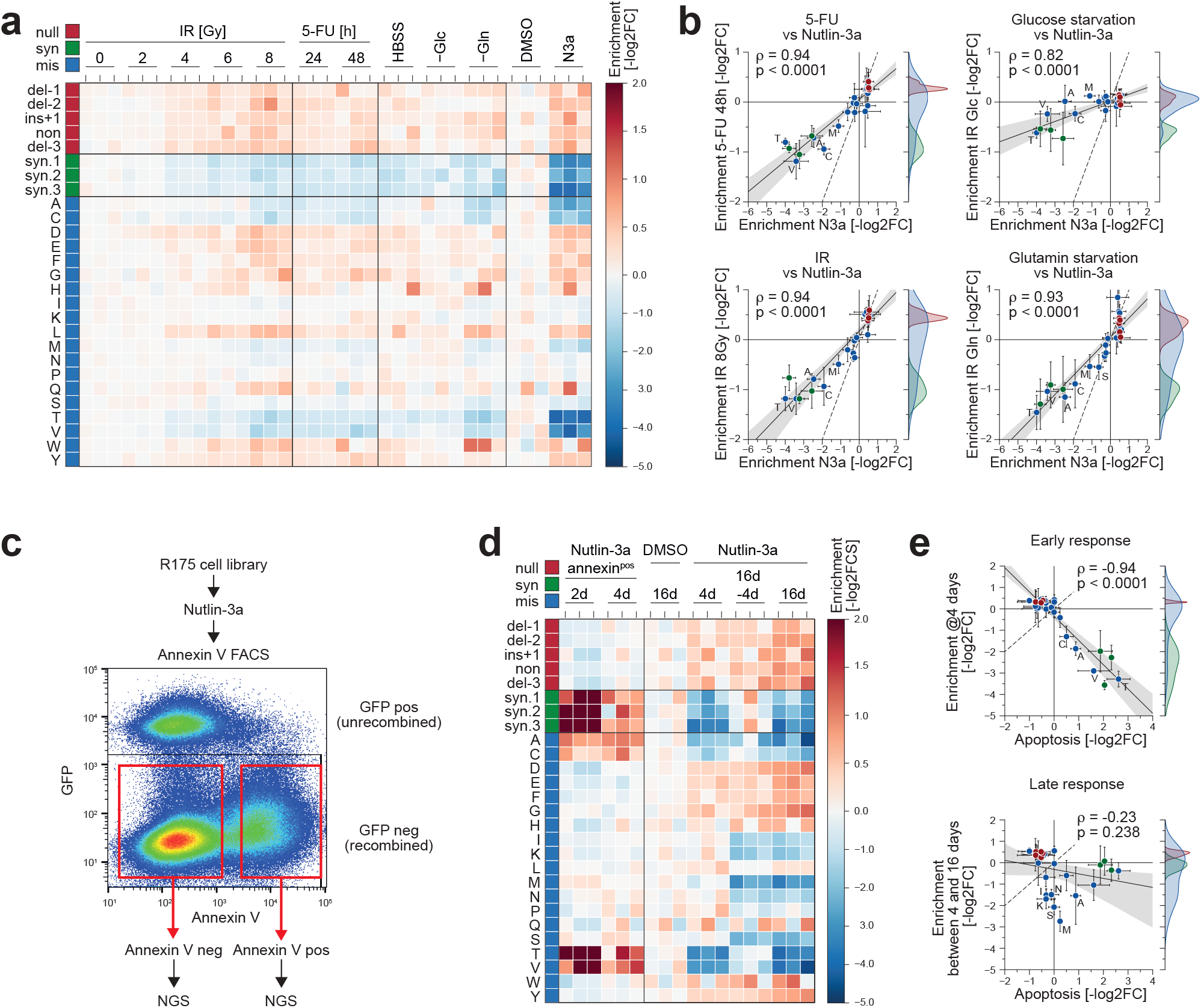
R175 variants: differential impact on stress responses and effector mechanisms. **a-b**, Comparison of different stress factors. **a**, Heatmap showing changes in variant abundance in response to DNA damage (IR, ionizing radiation; 5-FU, 5-fluorouracil) or nutrient starvation (HBSS, Hank’s buffered saline solution; -Glc, glucose starvation; -Gln, glutamine starvation) compared to control treatment with DMSO and N3a. Shown is the enrichment (n=3 biological replicates per condition) as the -log2 fold abundance change relative to the negative controls. **b**, Scatter plots illustrating the correlation between enrichment under DNA damage or nutrient deprivation and specific p53 activation with N3a. Shown is the mean ±SD enrichment (n=3 biological replicates) and Pearson correlation coefficient ρ. Dashed line, line of identity. **c-e**, Pro-apoptotic activity of R175 variants. **c**, Experimental scheme and a representative FACS scatter plot demonstrating the sorting strategy based on Annexin V staining. GFP-negative cells were gated to selectively analyze cells expressing the p53 variant, i.e., cells with successful deletion of the GFP-expressing LSL cassette after AV-Cre infection. **d**, Heatmap illustrating N3a-induced changes in variant abundance in the Annexin V-positive fraction (left) compared to the entire cell pool (right). Shown is the -log2 fold change (n=3 biological replicates) relative to the Annexin V-negative fraction (left) or DMSO-treated control cells (right). Lanes labelled as ‘16d-4d’ represent the difference between the 4d and 16d timepoint, reflecting late N3a-induced changes in variant abundance. **e**, Scatterplot showing the correlation between the early (4 days) and late (between 4 and 16 days) occurring N3a-induced changes in variant abundance versus their enrichment in the apoptotic cell fraction. Shown is the mean ±SD enrichment (n=3 biological replicates) relative to the DMSO-treated control and the Pearson correlation coefficient ρ. Dashed line, line of identity.

Moreover, we noted differences in the response kinetics of variants. R175A/C/T/V, much like the WT and syn variants, displayed an early depletion during the first 4 days, when N3a-induced apoptosis was most pronounced (Fig. 2e). In contrast, the depletion of R175I/K/M/N/S could only be observed at later time points, suggesting a different mechanism for their depletion compared to the rapid process of apoptosis. To test this hypothesis, we sorted and sequenced the apoptotic cells, positive for Annexin V, after 2 and 4 days of treatment (Fig. 3c). All syn and mis variants (R175A/C/T/V), which were rapidly depleted from the population, were strongly enriched in the apoptotic fraction (Fig. 3d,e), identifying apoptosis as the crucial mechanism in limiting tumor cell fitness. In contrast, the mutants that were depleted more slowly (R175I/K/M/N/S), were absent from the apoptotic fraction, supporting the idea that they are depleted through a mechanism independent of apoptosis, such as cell-cycle arrest or senescence. These findings underline the significant functional diversity among variants and highlight the power of CRISPR mutagenesis screens to uncover mechanistic differences.

### Saturating mutagenesis of the p53 DNA binding domain

We continued to screen more complex donor libraries, starting with one that included 210 non, syn and mis mutations targeting residues 176-185 that are involved in coordinating zinc and form the short helix H1 (Supplementary Fig. 4 and Supplementary Table 2). We then extended the screen to a comprehensive library that covered the p53 DNA-binding domain from exon 5 to 8 (amino acids 126 to 307) and encompassed approximately 94.5% of all cancer-associated missense mutations (Fig. 4a). The library consisted of 9,225 variants (Supplementary Table 3), including 1,919 single nucleotide substitutions (missense, nonsense or synonymous variants), which are the most common type of *TP53* mutations. Additionally, the library included 2,360 missense variants with single amino acid exchanges, 477 variants with additional nonsense mutations requiring 2 or 3 nucleotide substitutions, 2,560 variants with all possible single-nucleotide insertions, and 1,908 variants with 1, 2, and 3 bp deletions, including in-frame single amino acid deletions at each position.

**Fig. 4.**
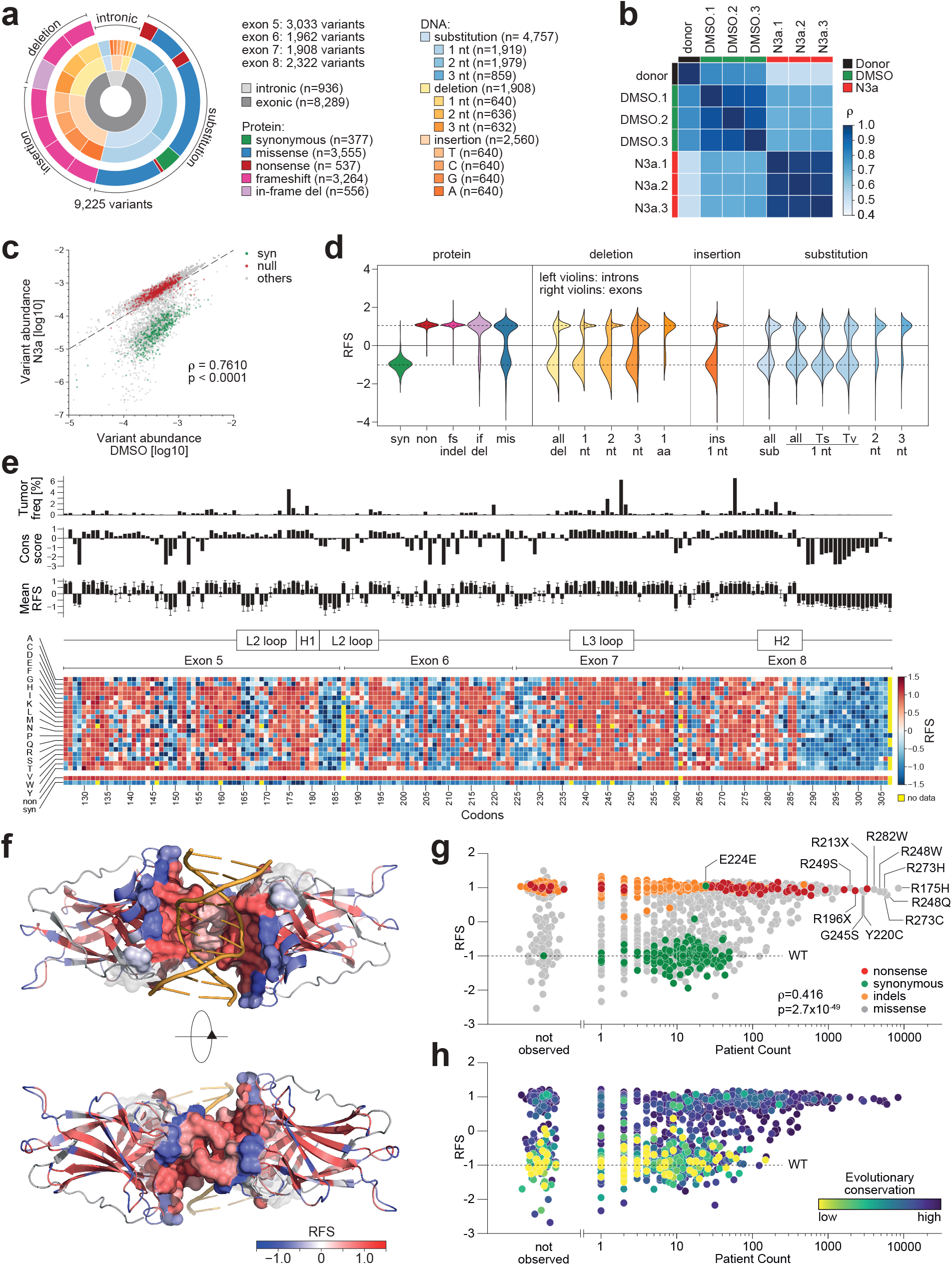
*TP53* DBD mutome screen. **a**, Composition of the *TP53* DBD mutagenesis library. **b-c**, Quality control plots. **b**, Heatmap showing pair-wise correlation coefficients (ρ, Spearman) between sample replicates. **c**, Scatter plot illustrating separation of variants under p53-activating N3a treatment. Shown is the median abundance of all variants under N3a versus DMSO treatment (n=3 biological replicates). Synonymous (syn) and nonsense (non) variants highlighted in green and red, respectively. ρ, Spearman correlation coefficient. Dashed line, line of identity. **d**, Distribution of relative fitness scores (RFS) for different variant classes. Left violin half shows distribution for intronic, right violin half for exonic variants. Fs, frameshift; if, in-frame; ins, insertion; del, deletion; indel, insertion or deletion; mis, missense; sub, substitution; Ts, transition; Tv, transversion; nt, nucleotide. **e**, Heatmap showing the RFS for all mis, syn, and non variants. Bar plots show for each codon the mutation frequency in the UMD *TP53* mutation database, the evolutionary conservation score, and the RFS (mean±SD) of all missense substitutions at this position. **f**, Structure of a DNA-bound p53 DBD dimer colored by RFS (Protein Data Bank 3KZ8^77^). The DBD-DNA and intra-dimer interaction interface within a distance of 10 Å is shown as a sphere model and superimposed on the cartoon model to highlight its sensitivity (red color, positive RFS values) to mutation. **g-h**, Scatter plots showing correlation between RFS and aggregated variant count in cancer patients listed in the UMD, IARC/NCI, TCGA, and GENIE databases. Variants are colored by the indicated mutation types (**g**) or evolutionary conservation (**h**). ρ, Spearman correlation coefficient.

To measure the abundance of individual variants in highly complex cell pools, an accurate calling of variant frequencies is required, close to the error rate of current deep sequencing technologies. Short-reads sequencing, such as Illumina’s sequencing-by-synthesis, currently provides the lowest error rates, but is restricted to a maximum length of 500-600 base pairs and therefore far too short to analyze genomic regions spanning multiple exons. To accommodate the length restrictions, the total library was subdivided into 4 sub-libraries, each covering one exon with 12 additional nucleotides of flanking intronic sequence. As described for the codon 175 library, each exon library was co-transfected with the *TP53*-targeting Cas9 nuclease into HCT116 LSL/Δ cells and edited cells were selected with puromycin and transduced with Cre to activate expression of the mutant. The complex cell pools were split and treated with either 10 µM N3a or DMSO solvent for 8 days. Genomic DNA was extracted, and the edited exon was selectively amplified by a nested PCR and analyzed by NGS (Supplementary Fig. 1a and Supplementary Table 4). Throughout the entire experiment, we maintained a mean coverage of at least 500 individually edited cells per variant. All screens were performed in three biological replicates for each exon and treatment, which correlated well within the treatment groups and with the original plasmid/donor library in the absence of N3a treatment (Fig. 4b and Supplementary Fig. 5a). Non mutations (LOF controls) and syn mutations (WT-like controls) did not significantly differ in their abundance, indicating highly efficient introduction of the donor library into the cells without detectable bias due to *TP53* status (Supplementary Fig. 5b). Following N3a treatment, the variant distribution changed substantially, reducing the correlation with the donor and DMSO-treated cell libraries (Fig. 4c and Supplementary Fig. 5a). Abundance of non and syn variants increased and decreased, respectively, while all other variants shifted from a uniform to a bimodal distribution, effectively separating LOF from WT-like variants (Supplementary Fig. 5b).

To normalize the enrichment scores of variants across multiple exons, the scores were transformed into Relative Fitness Scores (RFS), which scaled from -1 for the median of all synonymous variants to +1 for the median of all nonsense variants (Fig. 4d and Supplementary Fig. 5c,d). Frameshift-inducing variants, such as 1 or 2 bp exonic insertions and deletions (indels), were uniformly enriched, with RFS values similar to the nonsense controls (Fig. 4d). Interestingly, in-frame deletions of 3 consecutive bp also yielded mostly RFS values of +1, highlighting the sensitivity of the p53 DBD to even single amino acid deletions. The damaging effect of nucleotide substitutions was more variable, stronger for transversions versus transitions, and increased with the number of substituted nucleotides (Fig. 4d). Overall, 55.2% of all substitution variants showed a positive RFS value, indicating at least a partial functional impairment (Fig. 4d). The effects of intronic indels and substitution variants were less damaging and more variable, showing mostly negative RFS values, indicating preserved tumor suppressor activity (Fig. 4d).

Focusing on missense mutations, we explored the functional consequences of replacing each residue by every other amino acid (Fig. 4e). The screen returned reliable RFS values for 3,425 (99.05%) of all possible 3,458 missense mutant proteins, making this the most comprehensive study of the DBD mutome to date. The majority of missing variants mapped to exon boundary-spanning codons (G187, S261, A307) that could not be generated by substitutions within a single exon and were thus omitted during the library design phase. The effects of individual missense variants differed strongly based on their position and the type of amino acid substitution. Hierarchical clustering of missense mutants by RFS values differentiated codons according to their sensitivity to mutagenesis (Supplementary Fig. 5f). Codons that were highly susceptible to any amino acid substitution included the hotspots G245, R248, and R249, while the functional impairment of other hotspots such as R175 and R282 varied with the type of substitution (Supplementary Fig. 5e and f). Moreover, hydrophobic (V, I, L, M), aromatic (Y, F, W) and positively (H, R, K) or negatively (D, E) charged amino acids grouped together based on their biochemical functions, forming separate clusters (Supplementary Fig. 5f). This result confirms that substitutions with biochemically similar amino acids typically cause less damage than others, and that substitutions with structurally related amino acids result in comparable functional impairment.

The median RFS values of each codon were mapped onto the 3D protein structure, revealing a significant correlation between proximity to the DNA-binding surface and higher RFS values (Fig. 4f and Supplementary Fig. 5g). In addition, residues in the central hydrophobic core, critical for the thermal DBD stability, also showed significantly higher RFS values (Supplementary Fig. 5g-i). In contrast, solvent-exposed residues at the exterior of the DBD were more mutation-tolerant, with the notable exception of residues that directly contact DNA (R248) or are located at the inter-dimer interface (G199) (Supplementary Fig. 5h).

We next investigated whether the *in vitro* measured fitness effect of variants correlates with the prevalence in patient samples, based on over 150,000 *TP53* mutations reported in the UMD, IARC/NCI, TCGA and GENIE databases (Supplementary Table 5). The most frequent hotspot mutations as well as all other missense mutations with a patient count of more than 100 were strongly enriched and scored high RFS values (Fig. 4g, Supplementary Fig. 6a). Although far less prevalent than the most abundant missense mutations, also all of the screened nonsense and indel mutations reported in patient samples were enriched to positive RFS values. Confirming the specificity of the screen, all missense variants with negative (WT-like) RFS values showed lower patient counts. These WT-like missense variants have patient counts indistinguishable from syn mutations and benign polymorphisms^32^, and represent most likely passenger mutations. A notable outlier, the synonymous variant E224E, affects splicing and is described in more detail later. Comparing codon-level RFS values to evolutionary conservation scores from the ConSurf Database revealed a highly significant correlation (Fig. 4h, Supplementary Fig. 5j and Supplementary Table 6), indicating that p53 residues with high RFS scores are under evolutionary selection.

We also observed a high number of missense mutations at evolutionary conserved residues that scored high RFS values but were never or rarely reported in patients (Fig. 4g,h), raising the question whether these are false positives of our screen. Many of these variants were either 2 or 3 nt substitutions or single-nucleotide transversions (Supplementary Fig. 6b-d) that are all less frequent in cancer cells than single-nucleotide transitions^33^. While most of the hotspot mutations affect CpGs, none of the never observed high-RFS variants were at CpG sites. Nevertheless, irrespective of the type of mutation, variants with a positive RFS had significantly higher patient counts than those with negative RFS (Supplementary Fig. 6e-h), suggesting that they are selected for during tumorigenesis. We therefore also evaluated the sequence context of the screened variants for their mutational probability based on COSMIC mutational signatures that represent the major mutational processes accounting for base substitutions in cancer genomes (Supplementary Fig. 6i-l and Supplementary Table 7)^34^. Variants with positive RFS and high patient counts showed much higher mutational probabilities for all relevant mutational processes than high-RFS variants with low or zero patient counts (Supplementary Fig. 6i). In addition, variants with negative RFS despite high mutational probability have an average patient count below 100, underlining that these are random passenger mutations which are observed repeatedly in cancer patients only because of their mutational probability, not because they disrupt p53 function. When comparing variants with similar mutational probabilities, those with a positive RFS consistently showed significantly higher patient counts than those with a negative RFS (Supplementary Fig. 6k and l). A positive RFS therefore robustly identifies pathogenic loss-of-function variants, which are under positive selection during tumor development, and is therefore particularly informative for the pathogenicity classification of rare variants of unknown significance (VUS).

### Increased sensitivity of CRISPR screen for subtle loss-of-function

Previous studies have characterized the human *TP53* mutome by employing lentiviral overexpression of mutant cDNA libraries^4,8^. To compare the results of both types of screens, we transformed data from all screens to RFS values as described above, scaling the median of nonsense mutations to +1 and of synonymous mutations to -1 (Fig. 5a and Supplementary Table 8). Quality control plots showed the best separation between positive and negative controls and the highest statistical effect size for the CRISPR screen. The increased discriminatory power provided a clear separation of highly frequent cancer missense mutations from SNVs that were not reported as cancer-associated. In the cDNA overexpression screens, the RFS values for these functionally disparate variant types overlapped notably, possibly due to more variation in mutant expression levels from randomly genome-integrated lentiviral constructs.

**Fig. 5.**
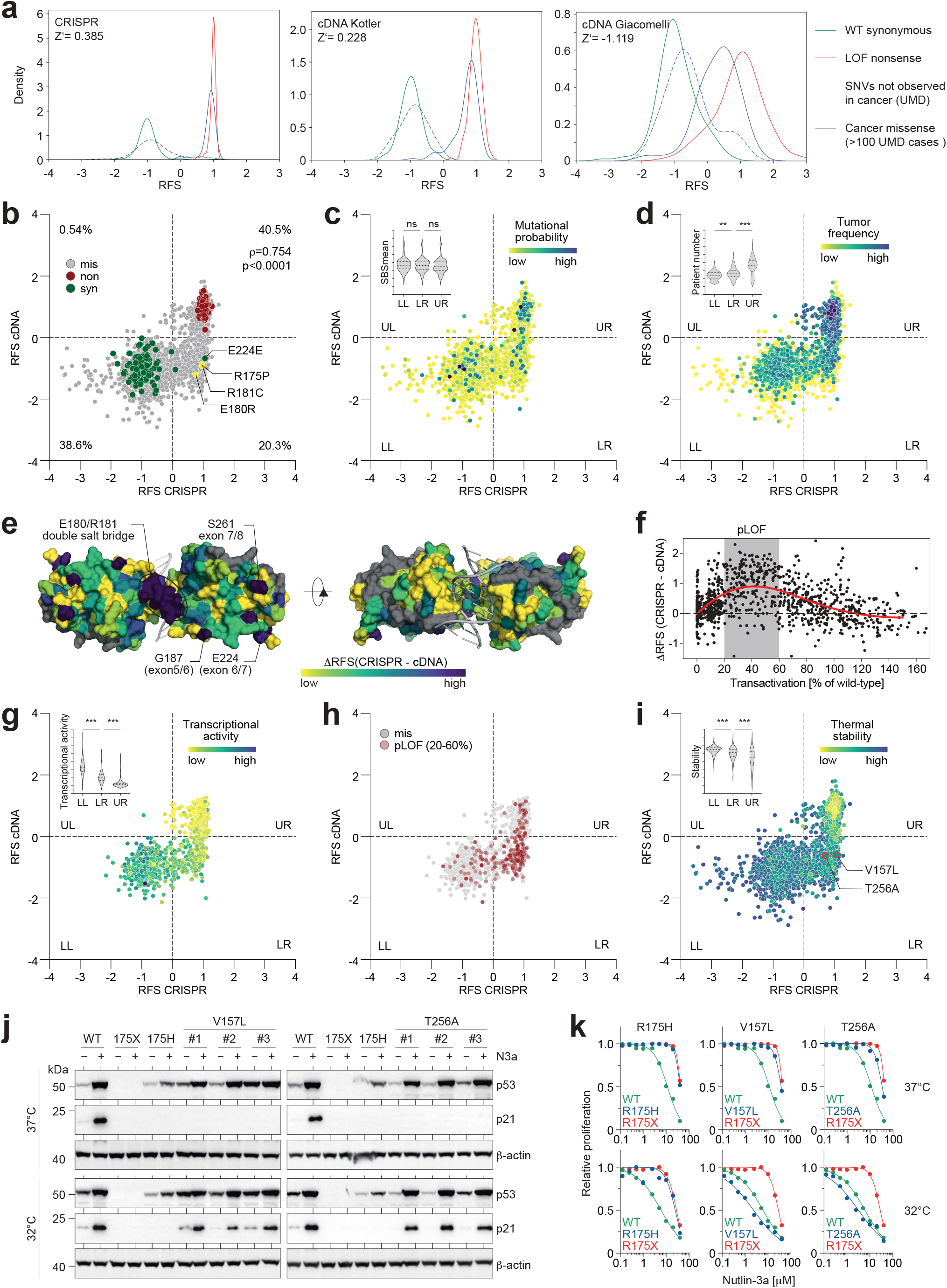
CRISPR screen reveals partial loss-of-function variants. **a**, Kernel density estimation (KDE) plots showing the distribution of RFS scores for the indicated groups of variants in the CRISPR versus cDNA-based mutome screens^4,8^. For comparison, all results from previously reported cDNA-screens were transformed to RFS by scaling the median of nonsense mutations to +1 and the median of synonymous mutations to -1. Z’ factors, a measure of statistical effect size, are stated as a quality parameter for the assay’s ability to separate positive (LOF nonsense) and negative (synonymous) controls. **b-d**, Scatter plots illustrating correlation between RFS values obtained by CRISPR mutagenesis and cDNA overexpression (Kotler et al., 2018^8^). Variants are categorized into 4 quadrants (LL, lower left; LR, lower right; UL, upper left; UR, upper right). Percentage of variants in each quadrant is given in **b**. Variants are colored by mutation type in **b**, average mutational probability (SBSmean) in **c**, or frequency in cancer patients in **d**. Inserted violin plots illustrate the value distribution in the three main quadrants. **e**, Structure of a DNA-bound p53 DBD dimer (Protein Data Bank 2AHI^76^) colored by the difference in RFS between the CRISPR and cDNA screen. Selected areas of high discrepancy are labelled. **f**, Scatter plot of the difference between CRISPR and cDNA screen versus the mean transcriptional activity of variants relative to wild-type p53 (WT) as measured in a yeast-based reporter system (Kato et al., 2003^7^). The area of 20-60% transcriptional activity (partial loss-of-function, pLOF) is shaded in grey; red line, cubic spline curve. **g-i**, Scatter plots showing correlation between RFS values obtained by CRISPR mutagenesis and cDNA overexpression. Variants are colored by transcriptional activity^7^ in **g**, classification as pLOF in **h**, or thermal stability as predicted by HoTMuSiC^39^ in **i**. V157L and T256A are highlighted in **i** with red outline and increased dot size. Inserted violin plots illustrate the value distribution in the three main quadrants. **j-k**, Temperature-sensitive function of LR variants V157L and T256A introduced into HCT116 LSL/Δ cells by CRISPR-HDR. **j**, Western blot of indicated HCT116 mut/Δ cell clones cultured with or without 10 µM N3a at 37 °C or 32 °C. **k**, Dose-response curves of N3a for representative cell clones, shown as mean relative proliferation (n=4 replicates), based on confluence measurements using real-time live-cell imaging. All violin plots show p values from one-way ANOVA and a post-hoc multiple comparisons test by Tukey (***, <0.001; **, < 0.01; ns, not significant). In all scatter plots, ρ denotes the Spearman correlation coefficient.

We first compared the results for individual missense mutants from the CRISPR screen with the study by Kotler^8^ (Fig. 5b). Both screens showed an overall good correlation with concordant classification as WT-like or LOF for all nonsense and synonymous variants and for ∼79% of the missense variants. Interestingly, 20.3% of missense variants were differentially classified as LOF by CRISPR and WT-like variants by cDNA screening (lower right (LR) quadrant variants). Hardly any variants (0.54%) were differentially classified in the other direction, indicating that the CRISPR screen has identified many more variants as potentially pathogenic than the cDNA screen. Even though LR variants had comparable mutational probabilities as all others (Fig. 5c, Supplementary Fig. 7a and b), they showed significantly higher patient counts than the WT-like variants in the lower left quadrant (Fig. 5d, Supplementary Fig. 7c), indicating that LR variants are positively selected during tumorigenesis and therefore likely pathogenic.

Importantly, three of the LR variants (R175P, R181C and E180R) are tumorigenic in mice with a (partial) LOF phenotype^35-37^, providing a further level of confirmation supporting the CRISPR classification of LR variants as pathogenic. In the 3D structure, a prominent region with discordant RFS values between CRISPR and cDNA screen mapped to the intra-dimer interface, where residues E180 and R181 form a double salt-bridge, mutations of which often result in partial LOF effects^38^ (Fig. 5e). Together this suggested that the CRISPR screen might be more sensitive for detecting subtle LOF phenotypes. Confirming this hypothesis, we observed the greatest differences between CRISPR and cDNA RFS values for variants with residual transcriptional activity, peaking in the 20-60% of wild-type range (Fig. 5f-h and Supplementary Table 8). An analysis of stability estimates by HoTMuSiC^39^ demonstrated a significantly higher thermal stability of LR versus UR (LOF) variants, but also a significantly lower stability compared to WT-like variants (Fig. 5i). All these observations were confirmed in a comparison with an independent cDNA screen by Giacomelli et al.^4^ (Supplementary Fig. 8).

To experimentally test the function of LR variants, we examined two representative mutations that are distant from functional interfaces, V157L and T256A, with total patient counts of 43 and 18 records, respectively. Both mutations clearly reduced the experimentally determined DBD melting temperature (V157L -5.7°C and T256A -3.7°C), but to a lesser extent than other more frequent mutations (V157F -8.5°C and Y220C -8.8°C) (Supplementary Table 9). When introduced into HCT116 LSL/Δ cells by CRISPR-HDR, both mutations rendered cells resistant to N3a comparable to the control mutations R175H and R175X (Fig. 5j and k). Given the modest degree of destabilization, we tested if the function is rescued at lower temperatures. Indeed, when cells were treated with N3a at 32°C, responsiveness to N3a, indicated by p21 expression and inhibition of cell proliferation, was rescued for both mutations, but not for R175H or R175X controls (Figure 5j and k).

Together, these results highlight that even a subtle loss of p53 function, resulting from a small degree of thermodynamic destabilization, can cause a profound increase in proliferative fitness. This increase was missed by conventional cDNA expression screens (where moderately increased unfolding rates may be offset by higher expression levels), but correctly identified when mutations are expressed at physiological levels from the endogenous gene locus. The CRISPR screen therefore revealed a novel set of pathogenic missense mutations with a low degree of thermal destabilization and possibly increased susceptibility to pharmacological rescue.

### Widespread splicing alterations and NMD

Interestingly, the largest differences between the CRISPR and cDNA screen map to a few isolated patches on the DBD surface formed by residues 187, 224, 225, and 261 (Fig. 5e), which are poorly conserved and highly tolerant to mutational perturbation in cDNA screens. Interestingly, these residues are all located at exon borders (Fig. 6a), suggesting that these mutations may affect RNA splicing. To assess the impact of individual variants on splicing, we sequenced cDNA from the cell libraries and correlated the abundance of all variants at the genome level with the abundance of correctly spliced cDNAs (Fig. 6b and Supplementary Table 10). Because alternative reading frames in the exons downstream of the DBD contain multiple stop codons, frameshift mutations in exons 5-8 result in premature termination of protein translation and are predicted to trigger nonsense-mediated decay (NMD), similar to nonsense mutations. As such, virtually all nonsense and frameshift mutations were significantly underrepresented at the mRNA level by ∼30-fold (Fig. 6c), which was sufficient to enhance proliferative fitness, as indicated by positive RFS values (Fig. 6d,e).

**Fig. 6.**
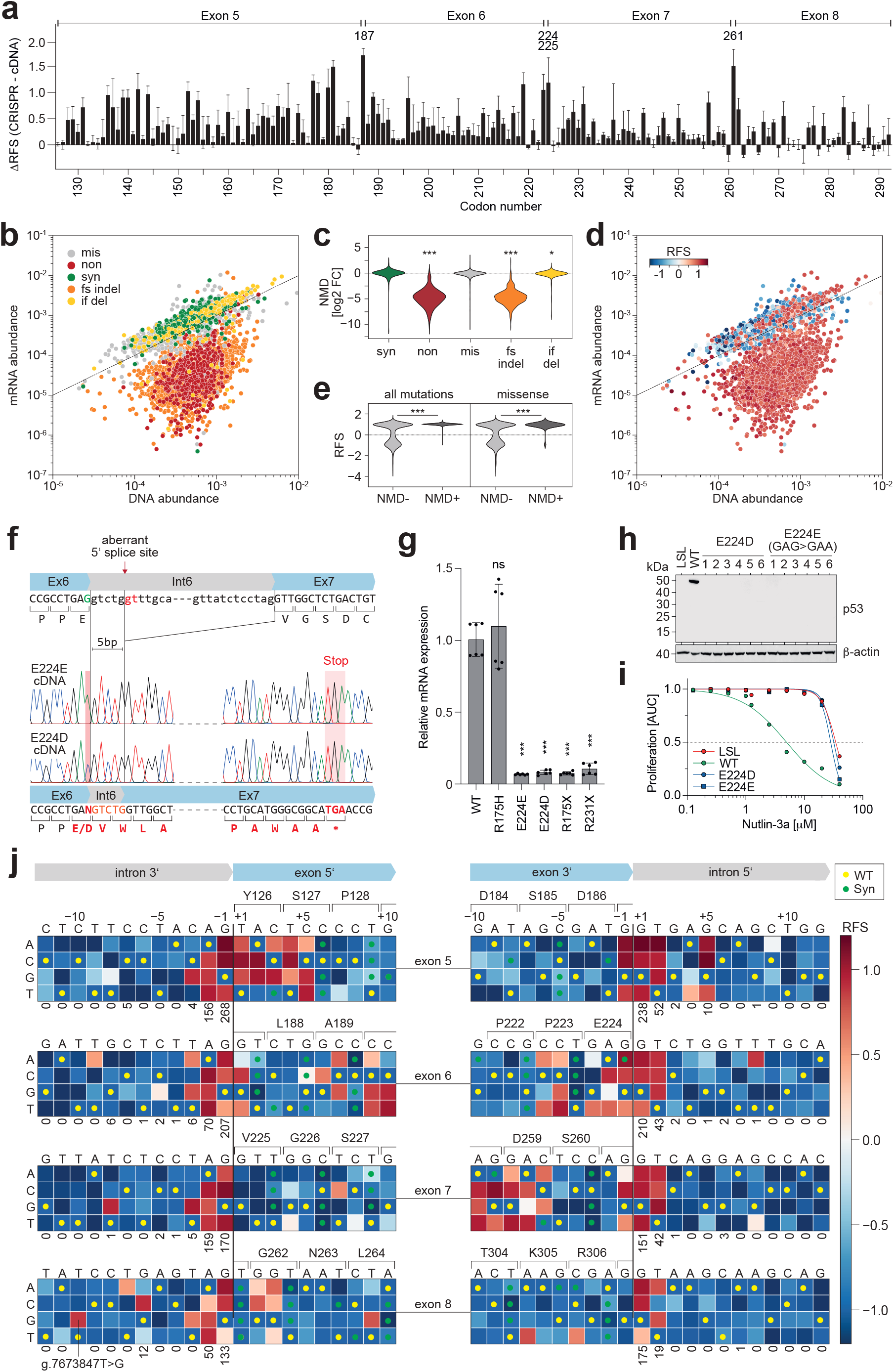
Splicing and nonsense-mediated decay (NMD). **a**, Barplot demonstrating high differences between CRISPR and cDNA screening results at exon borders (residues G187, E224, V225, and S261). Shown is the mean difference (±SD) of all missense variants at each codon. **b** and **d**, Scatter plots comparing the abundance of variants in the cell libraries at the level of genomic DNA and mRNA. Each dot represents the median abundance of a variant from n=3 biological replicates. Variants are colored by mutation type in **b**, and by RFS in **d**. Dashed line, line of identity. **c**, Violin plot showing NMD as the log2 fold-change in abundance at mRNA and DNA level by mutation type. ***, p<0.001; *, p<0.05; one-way ANOVA with multiple comparison by Tukey. **e**, Distribution of RFS values in variants (all or missense) according to NMD status. Variants with a log2 fold-change in abundance between mRNA and DNA <-2 were classified as NMD+. ***, p<0.001; two-sided Mann-Whitney test. **f-i**, Loss-of-function and NMD caused by E224E (synonymous) and E224D (missense) mutations. **f**, Aberrant mRNA splicing revealed by Sanger sequencing of cDNA. **g**, Quantitative reverse transcription PCR of indicated HCT116 mut/Δ cells. Shown is the *TP53* mRNA expression relative to WT as mean±SD (n=6 replicates). ***, p<0.001; one-way ANOVA with Dunnett’s multiple comparisons test. **h**, Western blot demonstrating lack of p53 protein expression in multiple HCT116 cell clones with E224/D mutations. **i**, Resistance of E224E/D clones to N3a. Proliferation was analyzed by real-time live cell imaging. Shown is the area under the proliferation curve relative to untreated. p53-null (LSL, red) and wild type (WT, green) are shown as reference. **j**, Impact of intronic and exonic variants near exon borders on tumor cell fitness. Heatmaps show the RFS of indicated single-nucleotide substitution variants. Wild-type and synonymous variants are indicated with yellow and green dots, respectively. The number of cancer patients in GENIE with mutations at exon-flanking intronic nucleotides is indicated below the heatmaps.

Interestingly, several missense mutations were also underrepresented at the mRNA level and associated with a LOF (Supplementary Fig. 9a). Most of these variants are located close to exon/intron borders (Supplementary Fig. 9b), supporting the hypothesis that they might affect mRNA splicing. While many are rare double or triple nucleotide substitutions, some are caused by single nucleotide substitutions (SNVs) and have been reported in several cancer patients (Supplementary Fig. 9c,d). Of particular significance are the G187, E224, and S261 mutations, which are prevalent in cancer samples and have been misclassified as WT-like in cDNA overexpression screens^4,7,8^. To study their functional impact in more detail, we introduced into HCT116 LSL/Δ cells the missense E224D (GAG>GAC) and synonymous E224E (GAG>GAA) mutations, which affect the last nucleotide of exon 5 and have been reported in 77 and 24 patients, respectively. Sequencing of cDNA revealed that both mutations attenuate the wild-type 5’ splice site (5’ ss) of intron 6 and enforce the use of an aberrant downstream 5’ss (Fig. 6f). The resulting inclusion of 5bp from intron 6 into the mature mRNA caused a frameshift and triggered a premature termination codon (PTC) present in an alternative reading frame of exon 7. As a consequence, the mRNA was subject to NMD, which prevented production of a truncated protein and rendered cells resistant to N3a (Fig. 6g-i).

We also noted mRNA depletion of one missense variant, L137Q (CTG>CAG), and one synonymous variant, G199G (GGA>GGT, not GGA>GGC or GGA>GGG), that were not located at exon/intron borders (Supplementary Fig. 9e and f). Closer inspection revealed that L137Q creates a new AG 3’ss in exon 5, whereas G199G (GGA>GGT) generates a cryptic GT 5’ss within exon 6. Both aberrant splice events were confirmed by sequencing reads spanning the aberrant splice junction. While the aberrant splice product caused by G199G (GGA>GGT) leads to a frameshift and NMD, the L137Q splice product is highly abundant and results in an in-frame deletion of 12 residues (Supplementary Fig. 9h). Although the G199G (GGA>GGT) transversion has a low mutational probability and has not yet been reported in a cancer patient, L137Q has been recurrently found in cancer patients and shows a LOF in the CRISPR screen that was not previously observed by cDNA screening.

Notably, 355 missense or synonymous SNVs created new exonic GT or AG dinucleotides that could theoretically function as cryptic splice sites. However, L137Q and G199G were the only SNVs not located at an exon/intron border that showed a more than 2-fold reduction of the expected full-length mRNA and resulted in a LOF (Supplementary Fig. 9a-d). Splice aberrations caused by intra-exonic SNVs are therefore far less frequent than predicted.

Apart from exonic splice mutations, we also observed LOF variants in the non-coding, exon-flanking, intronic regions that could be attributed to altered splicing. For example, we expectedly observed a LOF for most essential splice site mutations, i.e. for substitutions affecting the nearly invariant GT and AG dinucleotides at the intron ends (Fig. 6j). Moreover, we detected a deleterious impact of all SNVs at position 5 of intron 5, consistent with a strong prevalence for G at this position in the 5’ ss consensus sequence. While the impact of these mutations would have been predicted based on consensus sequences, introns 6-8 also harbor a G at this conserved position, but are tolerant to all substitutions. Consistently, the GENIE database lists 10 cases of various G5 substitutions for intron 5, but only a single case for introns 6-8. We also noted a LOF associated with the g.7673847T>G substitution in the 3’ region of intron 7 (Fig. 6j), which has been reported by the TCGA in a pancreatic adenocarcinoma patient^40^. Consistent with the creation of a cryptic 3’ss, we observed an aberrant inclusion of 9 bp of intronic sequence causing an in-frame insertion of 3 amino acids (Supplementary Fig. 9g,h). Different from cDNA-based screens, the CRISPR screen therefore discriminates pathogenic from non-pathogenic variants even in non-coding intronic regions.

### In vivo tumor growth selects for mutant p53 stabilization and metastatic behavior

Although p53 missense mutations have been described to promote tumorigenesis by a variety of GOF activities^12,13,17,18^, none of the missense mutations enhanced tumor cell fitness significantly beyond the beneficial effect of nonsense or frameshift mutations (Fig. 4e), suggesting that a proliferative GOF is not an intrinsic property of missense mutations. To examine if missense mutations can specifically enable the development of GOF properties, we compared the behavior of HCT116 cells with a missense and nonsense mutation when allowed to evolve *in vivo*. HCT116 R175H/Δ and HCT116 R175X/Δ cells were intravenously injected into mice, and after several weeks, established tumors were dissected from lungs and metastatic sites such as the liver. Explanted tumor cells were expanded *ex vivo* and re-injected into mice for a total of 3 mouse passages (Fig. 7a). In the course of months of serial *in vivo* propagation, we noted a progressive increase in migratory and invasive behavior of R175H cells from passage 0 (p0) to passage 3 (p3) that was not explained by differences in proliferation and not observed with the R175 nonsense mutation (Fig. 7c-j, Supplementary Fig. 10a-d). Migration and invasion of R175H-p3 cells was significantly reduced by knock-down or knock-out of the mutant (Fig. 7e-j, Supplementary Fig. 10c-h). Moreover, knockout of the mutant in R175H-p3 cells had no impact on the growth of subcutaneously grown tumors, but significantly impaired metastasis to the liver (Fig. 7k-m). Thus, R175H-p3 cells do not become addicted to R175H, but exhibit metastatic properties that depend on the continued presence of the mutant and are therefore most likely mediated by the mutant protein.

**Fig. 7.**
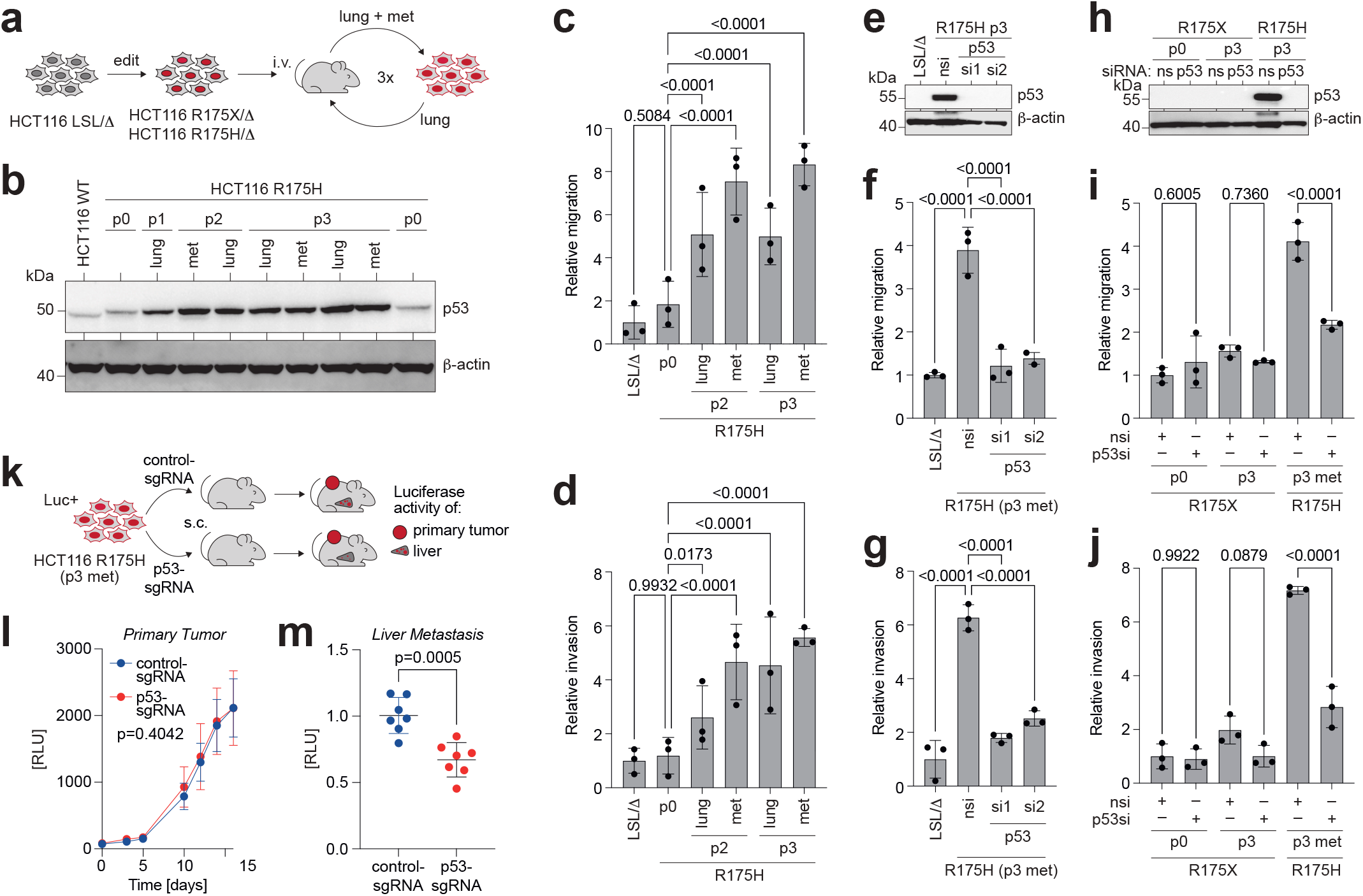
R175H enables evolutionary development of pro-metastatic properties. **a**, *In vivo* tumor evolution model. HCT116 cells with R175H and R175X mutations were grown in mice after intravenous injection. Tumors from lungs and metastatic sites were explanted, expanded in cell culture and re-injected for up to 3 mouse passages. **b**, Western blot of R175H-mutant HCT116 cells showing protein stabilization following multiple rounds of mouse passaging (p0-3). The R175H mutant is expressed as the 72P polymorphic variant, resulting in an apparent higher molecular weight compared to the wild-type protein. **c-d**, Transwell assays for migration and invasion after indicated rounds of mouse passaging demonstrate gain of pro-metastatic activity. Shown is the mean ±SD (n=3 experiments with 3 replicates each) relative to original p53-null (LSL/Δ) HCT116 cells and results from two-way ANOVA with Dunnett’s multiple comparisons test. **e-j**, Transwell migration and invasion is dependent on R175H protein expression. nsi, non-silencing siRNA; p53si, p53-targeting siRNA. Shown is the mean ±SD (n=3 replicates) and results from one-way ANOVA with Sidak’s multiple comparisons test. Western blots in **e** and **h** confirm lack of p53 protein expression in R175X cells and efficient siRNA-mediated depletion of R175H. **k-m**, *In vivo* metastasis assay. **k,** HCT116 R175H/Δ p3 cells were dual-labelled with firefly and secreted Gaussia luciferase, transfected with CRISPR nucleases (control or targeting p53), and subcutaneously injected into immunodeficient mice. **l**, Primary subcutaneous tumor growth was measured based on secreted Gaussia luciferase levels in blood samples. Shown is the mean ±SD (n=7 mice per group). p-value of group factor from two-way ANOVA. **m**, Metastasis to liver was measured based on firefly luciferase activity in whole liver homogenate. Shown is the mean ±SD luciferase activity relative to control-sgRNA (n=7 mice per group). RLU, relative light units; p-value from two-sided unpaired t-test.

Mutant p53 is inherently unstable in non-transformed cells and GOF activities require its stabilization^41^. This has been mechanistically linked to the malignant transformation process, during which oncogenes and p53 loss of heterozygosity upregulate heat shock proteins which shield mutant p53 from proteasomal degradation^42,43^. Even though HCT116 cells are transformed cells derived from a clinically manifest colorectal cancer, CRISPR mutagenesis of the *TP53* gene locus has resulted in HCT116 mut/Δ cells with mutant protein levels that were similar to wild-type p53, and remained under tight Mdm2 control (Fig. 1c). This demonstrates that p53 missense mutants are inherently unstable not only in non-transformed but also in fully transformed cells, unless specific alterations stabilize the mutant protein. Intriguingly, mutant p53 protein levels in HCT116 cells increased progressively upon serial *in vivo* passaging (Fig. 7b), suggesting that *in vivo* tumor growth selects for cells which have acquired mechanisms to stabilize the mutant protein. Although metastatic properties of R175H-p3 cells were diminished by mutant p53 depletion, mutant p53 stabilization by N3a did not induce metastatic behavior in R175H-p0 cells (Supplementary Fig. 10i-k), demonstrating that stabilization is required but not sufficient for metastatic behavior.

## Discussion

The comprehensive CRISPR screen reported in this paper covers approximately 94.5% of all cancer-associated *TP53* mutations. This study should be viewed in the context of previous high-throughput screens of the *TP53* mutome using overexpression of mutant cDNA in yeast or human cell lines^4,7,8^. The CRISPR approach showed superior separation of WT-like and LOF variants due to improved precision and reduced variability in calling fitness effects (Fig. 5a). Despite the fact that lentiviral integration is not entirely random, the increased variability in mutant expression levels in the cDNA overexpression screens may be explained by highly variable lentiviral integration, occurring at differently active genomic regions^44^. In contrast, CRISPR introduces the mutation into a defined genomic locus, here the endogenous *TP53* gene, resulting in highly consistent protein expression and reproducible fitness effects upon N3a treatment between independently edited clones (Supplementary Fig. 1f-h).

Due to the observed improvement in discriminatory power, we identified approximately 20% of the missense variants as fitness-promoting mutations that were previously missed by cDNA-screening (Fig. 5b). A closer examination showed that these variants have a similar mutational probability but are significantly more frequent in cancer patients compared to synonymous or WT-like variants (Fig. 5c and d), indicating that they are pathogenic mutations selected during tumorigenesis, which is further demonstrated by the tumorigenicity of some of these variants in mice^35-37^. This suggests that the pathogenic nature of many *TP53* variants has been underestimated previously due to technical limitations that can be overcome by a CRISPR screening approach.

Interestingly, many of these variants were predicted to be thermally destabilized by only a few degrees of temperature (Fig. 5i), significantly less than some of the more frequent structural hotspot mutants such as Y220C^45^. This could explain their overall higher residual transcriptional activity (Fig. 5g and h) and their lower frequency in cancer patients (Fig. 5d), but also identifies them as potentially temperature-sensitive, which was confirmed by a detailed analysis of representative mutations (V157L and T256A, Fig. 5j-k). From a translational perspective this suggests that the folded, active conformation of these mutants might be more easily rescued^24,46^, making these variants an especially interesting subset for pharmacological targeting approaches. It is tempting to speculate that cancer patients with variants from this subset might benefit most from therapeutic approaches suggested for temperature-sensitive mutations, such as hypothermia and antiparasitic antimonials^47,48^.

Splicing effects, for which cDNA-based screens are obviously blind, also account for some of the fitness-promoting mutations in our screen. As expected, substitution variants that affect regular splicing tend to cluster at exon/intron borders, either by altering the GT and AG dinucleotides (5’ and 3’ss) at the intron ends or by changing the first or last exonic nucleotides, where mutations can weaken the regular splice sites and promote the usage of alternative ones^49^. We observed two intra-exonic substitutions (L137Q missense and G199G (GGA>GGT) synonymous) that created cryptic splice sites, resulting in aberrantly spliced transcripts that promote tumor cell fitness (Supplementary Fig. 9). While the L137Q variant caused an in-frame deletion and was readily detectable at the mRNA level, the G199G mutation led to a frameshift and triggered NMD. Notably, these were the only two SNVs outside the exon/intron border regions that modified splicing patterns. This is especially noteworthy because many more SNVs create potential cryptic splice sites but do not appear to disrupt regular splicing or impair p53 function. It was anticipated that SNVs located at putative exonic splicing enhancer/silencers would have splicing effects. However, the lack of splice alterations by most exonic SNVs suggests that *TP53* splicing in the DBD region is robust against SNVs. This means that single-nucleotide alterations are typically not enough to establish a functional splice site or to interfere with the function of splicing enhancers/silencers. However, SNVs that create a novel splice site are removed from the mRNA during splicing, so we can only infer their splicing effect indirectly from the absence of a normally spliced mRNA encoding the variant. It is therefore possible that some SNVs produce a mix of normal and additional, aberrantly spliced transcripts. To fully understand the extent and functional impact of such variants, a comprehensive analysis is needed that combines DNA and mRNA analysis of mutome libraries at the single-cell level.

Our study primarily used Mdm2 inhibition with N3a as a specific method to activate p53. This approach is based on the understanding that p53 is ultimately activated by disrupting the p53-Mdm2 interaction, regardless of the stress factor involved. As such, we observed a similar variant enrichment/depletion pattern with various p53-activating stress factors, including DNA damage and metabolic stress, compared to N3a or other Mdm2/Mdmx inhibitors (Fig. 3a and b). However, the level of enrichment/depletion was weaker with other stressors as additional stress pathways not related to p53 also impact on tumor cell fitness. Interestingly, the study noted that loss of p53 function protected from metabolic stress, even though wild-type p53-mediated cell-cycle arrest through p21 induction was shown to promote survival under conditions of temporary nutrient starvation^50,51^. The difference in our results might be due to different treatment protocols, including our use of harsh starvation conditions that aimed at killing most cells. In addition, the donor vectors used to produce the mutant libraries encode proline at the polymorphic codon 72, which is a weaker activator of p21 and protects less from nutrient deprivation compared to the arginine variant (R72) found in the parental HCT116 cells^52^.

It is widely believed that p53 is more frequently hit by missense mutations compared with other tumor suppressors because they can provide the mutant protein with pro-tumorigenic GOF properties^12,13^. Our study found no evidence of a fitness advantage from missense mutations compared with null mutations (Fig. 4). This may be due to the fact that the stabilizing effect required for GOF properties was not observed in the p53-mutant HCT116 cells. All of the missense mutants showed normal expression levels and only accumulated after N3a treatment (Fig. 1c), which indicates proteasomal degradation by Mdm2 under normal conditions. The stabilization of p53 mutants in tumors is usually achieved through oncogenic alterations that increase the levels of heat shock chaperones, which protect the mutant protein from degradation^41,53^. This requires the loss of the remaining normal allele^43,54^ as wild-type p53 represses the expression of heat shock transcription factor 1^42^. HCT116 cells are fully malignant cells derived from a patient with colon cancer, and the mutant derivatives generated in our study do not express normal p53 (Fig. 1c). Neither malignancy nor loss of heterozygosity is therefore sufficient to stabilize the mutant protein. Nonetheless, HCT116 R175H cells showed stabilization of the mutant protein and acquired pro-metastatic GOF properties when serially propagated *in vivo* (Fig. 7). This suggests that a subset of R175H-mutant cells can acquire mechanisms to stabilize the mutant protein and develop GOF properties. Testing this on other mutants would be valuable. Unfortunately, conducting a comprehensive *in vivo* screen with the available mutant cell libraries is currently unfeasible because the limited tumor-initiating potential of HCT116 cells makes it difficult to maintain a sufficient coverage. A cell-based screen in a cell line with mutant-p53 driven pro-metastatic properties could be an alternative but might face gene editing challenges due to the genetic instability of p53-mutated cells.

## Limitations

The study was limited in its ability to examine the dominant-negative activity of variants that promotes tumorigenesis in the presence of a remaining wild-type allele^55,56^. Our HCT116 LSL/Δ screening cell line has an inactive second allele. To study dominant-negative effects, we attempted to generate an LSL/WT cell line, but gene editing in these cells was less efficient, potentially due to wild-type p53 interference with HDR-mediated DSB repair^21-23^. Additionally, these cells were genetically unstable and, like other p53+/- cells^54,57^, underwent LOH at high frequency. We also confirmed our results in a different cellular context, the p53 wild-type, non-small cell lung cancer cell line H460. We successfully introduced single mutations and a small R175 library into this cell line using the same approach as in HCT116 (Supplementary Fig. 3). The results were similar to those with HCT116 cells, indicating similar effects of variants regardless of cell type. However, the transfection and editing efficiency were inadequate for larger screens. HCT116 cells have already been widely used in the pre-CRISPR era for mutagenesis by homology-directed repair due to their mismatch repair deficiency^58-60^, and dominant-negative inhibition of MMR has also been shown to improve gene editing with prime editors^61^. It is possible that the exceptional HDR efficiency we observed in our screen was facilitated by this, and that transient inhibition of MMR might be a strategy to enable similar screens in other model systems.

## Conclusion

In summary, the CRISPR screen of the *TP53* cancer mutome provides a comprehensive functional annotation of its impact on tumor cell fitness. The high-resolution analysis identified numerous variants with subtle loss-of-function phenotypes that are selected during tumorigenesis and therefore should be considered pathogenic in genetic counseling or personalized cancer medicine. The temperature-sensitive phenotype of such variants suggests they may be especially susceptible to pharmacological rescue approaches. With the exception of SNVs located at exon/intron borders, few variants affected splicing, demonstrating the robustness of the *TP53* splicing pattern. Notably, the study found no evidence of a fitness advantage from missense mutations compared with null mutations, and neither malignancy nor loss of heterozygosity were sufficient to stabilize the mutant protein, underlining that stabilization and GOF effects are not intrinsic properties of missense mutations but arise during tumor evolution in a mutant p53-dependent manner.

## Methods

### Cell Culture

The human colorectal carcinoma cell line HCT116 and human embryonic kidney cell line HEK 293T were obtained from the American Tissue Collection Center (ATCC, CCL-247) and cultured in Dulbecco’s modified Eagle’s medium (DMEM, Gibco, 41966-029) supplemented with 10% heat-inactivated fetal bovine serum (FBS, Sigma-Aldrich, S0615) and 100 U/ml penicillin and 100 µg/ml streptomycin (Gibco, 15140122). The human large-cell lung cancer cell line NCI-H460 (H460) was obtained from ATCC (HTB-177) and cultured in Roswell Park Memorial Institute (RPMI) 1640 medium (Gibco, 61870) supplemented with 10% FBS and 100 U/ml penicillin and 100 µg/ml streptomycin. Cell lines were cultivated in a humidified atmosphere at 37 °C and 5% CO_2_ and detached using 0.05% Trypsin-EDTA (Gibco, 15400054).

### Design of mutome library

The wild-type *TP53* sequence was derived from the human Ensembl genome, revision 96 (GRCh38). As the cloning procedure uses BbsI-mediated Golden Gate Cloning, BbsI recognition sites present within *TP53* exon 4 to intron 9 were silently mutated and the resulting sequence was used as a template for library generation. Using transcript ENST00000269305 (RefSeq NM_000546), the sequences of exons 5, 6, 7 and 8, including 12 nucleotides of flanking intron sequence, were selected and subjected to *in silico* mutagenesis. 13 nucleotides of the introns flanking this ’mutatable’ sequence were added so that this variable region was framed by short constant regions that would remain the same to all resulting synthetic oligonucleotides. To generate an exhaustive set of ’mutated’ oligonucleotides that deviate from the wild-type sequence by a single mutation, the variable region was altered in the following way. Initially, each nucleotide was substituted with every other nucleotide, resulting in a comprehensive set of all single-nucleotide variants (SNVs). To include amino acid substitutions that cannot be achieved by a single-nucleotide substitution, we added double-nucleotide variants (DNVs) and triple-nucleotide variants (TNV) to generate each possible amino acid substitution and non-sense mutation. In the case of multiple possible codon exchanges, we prioritized the change with the smallest hamming distance to the reference. To account for insertions, each possible single nucleotide was inserted at every position of the variable region, including the intronic region, resulting in a set of all possible insertions of size 1 bp. Finally, a deletion set was generated by deleting up to three nucleotides at every position of the variable region, thus creating a set of all possible deletions of size one to three.

### Generation of plasmids for CRISPR mutagenesis by homology directed repair (HDR)

Homology arm 1 ranging from exon 4 to intron 4 (chr17:7,675,788-7,676,168) of the *TP53* gene was PCR-amplified from genomic DNA of HCT116 cells using the following primers: HA1_BsrGI_fw 5’-CAC CTA TAT GTA CAA GAG GCT GCT CCC CCC GTG-3’, HA1_BsaI_rev 5’-TAT AGA GAC CGA TGG ATA AAA GCC CAA ATT C-3’ and cloned into the multiple cloning site 1 (MCS1) of the vector MCS1-EF1α-GFP-T2A-Puro-pA-MCS2-PGK-hsvTK (HR700, System Biosciences) using BsrGI (New England Biolabs, R3575) and BsaI (New England Biolabs, R3733). Homology arms 2 ranged from intron 4 to intron 6 (for mutagenesis of exons 5 and 6, chr17:7,674,377-7,675,787) or from intron 4 to intron 9 (for mutagenesis of exons 7 and 8, chr17:7,673,145-7,675,787) with BbsI-recognition sites flanking the region to be mutated (R175: chr17:7,675,059 and 7,675,088; H1 helix: chr17:7,675,059 and 7,675,085; Ex5: chr17:7,675,036 and 7,675,254; Ex6: chr17:7,674,842 and 7,674,989; Ex7: chr17:7,674,164 and 7,674,308; Ex8: chr17:7,673,684 and 7,673,855) were purchased as custom gene synthesis (GeneArt, Thermo Fisher) and cloned into MCS2 of HR700 using MluI (New England Biolabs, R3198) and SalI (New England Biolabs, R3138). In total, we generated 6 different HR700 donor vectors for cloning of libraries targeting R175, codon 176-185 (short: H1 helix), exon 5, 6, 7, and 8.

For generation of R175 and H1 helix plasmid libraries, complementary single-stranded oligonucleotides containing the desired mutations were purchased (Eurofins Genomics) and annealed individually to generate double-stranded DNA containing suitable overhangs. Double-stranded oligonucleotides were purified using PCR purification kit (QIAGEN, 28106) and cloned into HR700 vectors using BbsI-mediated Golden Gate Cloning.

For generation of exon-wide plasmid libraries, single-stranded oligonucleotide pools containing the desired mutations were purchased (oPools, Integrated DNA Technologies) and BbsI-recognition sites were introduced by PCR-amplification ensuring a coverage of 1 x 10^6^ for each mutation using the following primers: Exon 5 fw 5’-CAA TAT GAA GAC CTC TGT CTC CTT C-3’, Exon 5 rev 5’-ATA TAT GAA GAC CGT CTC TCC AGC C-3’, Exon 6 fw 5’-CAA TAT GAA GAC CTG ATT CCT CAC T-3’, Exon 6 rev 5’-ATA TAT GAA GAC CAG AGA CCC CAG T-3’, Exon 7 fw 5’-CAA TAT GAA GAC ATC TTG GGC CTG T-3’, Exon 7 rev 5’-ATA TAT GAA GAC TGC AGG GTG GCA A-3’, Exon 8 fw 5’-CAA TAT GAA GAC GCT TCT CTT TTC C-3’, Exon 8 rev 5’-ATA TAT GAA GAC ACC GCT TCT TGT C-3’. Amplified oligos were purified using PCR purification kit (QIAGEN, 28106) and cloned into HR700 vectors using BbsI-mediated Golden Gate Cloning.

Plasmid libraries were transformed into MegaX DH10B T1R Electrocomp E. coli (Invitrogen, C640003) and seeded on 2 (R175), 10 (H1 helix) or 30 (exon-wide libraries) 15 cm agar plates containing 50 µg/ml kanamycin (Carl Roth GmbH, C640003). After 16 h growth at 37 °C, colonies were scraped off, pooled into 100 ml (R175), 400 ml (H1 helix) or 1.2 l LB-medium and incubated for 4 h at 37 °C before extracting plasmid DNA using Nucleobond Xtra Midi kit (Macherey-Nagel, C640003) according to the manufacturer’s protocol.

Donor HDR plasmids for single mutations were either generated using annealed or PCR-amplified oligonucleotides as described above. Correctness and integrity of plasmids was validated using Sanger sequencing (LGC Genomics) or next-generation sequencing (NGS). Plasmids for delivery of Cas9 and sgRNAs were generated using BbsI-mediated Golden Gate cloning of annealed single-stranded oligonucleotides into pX330-U6-Chimeric_BB-CBh-hSpCas9 (pX330, Addgene #42230), pSpCas9(BB)-2A-Puro (PX459) V2.0 (pX459_puro, Addgene #62988), pSpCas9(BB)-2A-Hygro (pX459_hygro, Addgene #127763) or pSpCas9(BB)-2A-Blast (pX459_blast, Addgene #118055).

### Generation of HCT116 LSL/Δ and NCI-H460 LSL/Δ/Δ cells

2.5 x 10^4^ HCT116 cells were transfected with 1.25 µg pX330_sgIn5 (sgRNA 5’-TCA GTG AGG AAT CAG AGG CC-3’) and 1.25 µg HR700, which contained wild-type homology arms 1 and 2 flanking the LSL cassette, using Lipofectamine 2000 (Thermo Fisher Scientific, 11668019) according to the manufacturer’s protocol. Cells were selected with 1 µg/ml puromycin (Invivogen, ant-pr) and single cell clones were isolated. Single cell clones were chosen based on Nutlin-3a (N3a) and puromycin resistance and analyzed by PCR for a Δ-allele with an inactivating deletion in intron 5 and a second allele containing the LSL-cassette (HCT116 LSL/Δin5). The absence of the LSL-cassette on the Δ-allele was confirmed using primers: Intron4_fw: 5’-CCC TTT GGC TTC CTG TCA GTG-3’, Exon7_rev: 5’- GAT GGT GGT ACA GTC AGA GCC-3’. Sanger sequencing showed deletion of chr17:7,674,986 – 7,675,001. The presence of the LSL-cassette was validated with two PCRs, one spanning the upstream end (Intron1_fw: 5’-GGT GAC CCA GGG TTG GAA GTG T-3’, GFP_rev 5’- TGG GGT GGA TGG CGC TCT TGA A-3’) and the other spanning the downstream end (LoxP_fw 5’- GGG GGC TGT CCC TAG ATC TAT AA-3’, Exon7_rev 5’-GAT GGT GGT ACA GTC AGA GCC-3’). Finally, digital PCR for GFP (TaqMan Copy Number Assay, Applied Biosystems, 4400291) was performed using QuantStudio 3D Digital PCR 20K Chip V2 (Applied Biosystems, A26316) to confirm the presence of only a single copy of the LSL-cassette in the genome. The respective single cell clone of HCT116 LSL/Δin5 was then transfected with pX330_sgPuro (sgRNA 5’-CAC GCC GGA GAG CGT CGA AG-3’) to knock out the *pac* gene present in the LSL cassette. After validation of the puromycin sensitivity, HCT116 LSL/Δin5 cells were further transfected with pX459_hygro_sgIn7 (sgRNA 5’-CCA CTC AGT TTT CTT TTC TC-3’) to generate HCT116 LSL/Δin5+7 cells for mutagenesis of exon 7 and 8. After selection with 250 µg/ml hygromycin (Invivogen, ant-hg), single cell clones were screened via PCR (Δin5+7 allele: Intron4_fw 5’-CCC TTT GGC TTC CTG TCA GTG-3’, Exon8_fw 5’-AGG CAT AAC TGC ACC CTT GG-3’; LSL-allele: LoxP_fw 5’-GGG GGC TGT CCC TAG ATC TAT AA-3’, Exon8_fw 5’-AGG CAT AAC TGC ACC CTT GG-3’). A respective single cell clone of HCT116 LSL/Δin5+7 with distinguishable deletions (LSL-allele: chr17:7,673,970-7,673,995; Δ-allele: chr17:7,673,986-7,674,259) on both alleles was chosen for further experiments.

H460 LSL/Δ/Δ cells were generated from NCI-H460 using the same procedure, with special attention given to the fact that this cell line has three *TP53* alleles, meaning it must contain two Δ alleles and one LSL allele.

### Generation and treatment of single mutants and cell libraries

For generation of single mutants, 2.5 x 10^4^ HCT116 LSL/ΔIn5, HCT116 LSL/ΔIn5+7 or H460 LSL/Δ/Δ cells were transfected with 1.25 µg LSL allele-specific sgRNAs (pX459_blast_In5^LSL^, sgRNA 5’-GTG AGG AAT CAG AGG ACC TG-3’ or pX459_blast_In7^LSL^, sgRNA 5’-CTT TGG GAC CTA CCT GGA GC-3’) and 1.25 µg of the corresponding HR700 vector carrying the intended mutation using Lipofectamine 2000 according to manufacturer’s protocol. Transfected cells were selected with 20 µg/ml blasticidin (Invivogen, ant-bl) for 3 days and 1 µg/ml puromycin for 7 days, before single cell clones were isolated and the presence of the mutation was validated through edit-specific PCR and Sanger sequencing. Finally, cells were infected with AV-Cre (ViraQuest Inc., Ad-CMV-Cre, MOI20 for HCT116 cells, MOI250 for H460 cells) and expression of the mutant was confirmed via cDNA sequencing and Western blot analysis.

For the generation of R175- and H1 helix libraries, 4 x 10^6^ or 12 x 10^6^ HCT116 LSL/ΔIn5 cells were transfected with 6.25 µg or 18.75 µg pX459_blast_In5^LSL^ and 6.25 µg or 18.75 µg of the HR700 vector library, respectively, using Lipofectamine 2000. For the generation of R175-libraries in H460 cells, 4 x 10^6^ H460 LSL/Δ/Δ cells were transfected with 20 µg pX459_blast_In5^LSL^ and 20 µg HR700 vector library using the Neon Transfection System (Thermo Fisher Scientific, MPK10025). Transfected cells were selected for 3 days with 20 µg/ml blasticidin and for 7 days with 1 µg/ml puromycin. 8 x 10^6^ (R175) or 1.6 x 10^7^ (H1 helix) cells were then infected with AV-Cre and, after 5 days, the cell library was divided and treated with 10 µM N3a (BOC Sciences, B0084-425358), 75 nM RG7388 (MedChemExpress, HY-15676), 10 µM RO-5963 (Calbiochem, 444153), 1 µM MI-773 (Selleckchem, S7649), 750 nM AMG 232 (MedChemExpress, HY-12296) or the respective volume of DMSO (Carl Roth GmbH, 4720) as solvent control for 8 days. For irradiation experiments, an X-RAD 320iX tube was used with settings of 320 kV voltage and a current of 8 mA, with a dose rate ∼1 Gy/min. Cells were further cultivated for 8 days after irradiation. 5-Fluorouracil (pharmacy of the Universitätsklinikum Gießen and Marburg) was administered at a concentration 5 µM for 24 h or 48 h, and cells were further cultivated for 8 days after treatment. For mutant p53 reactivation studies, cell libraries were treated with either 12.5 µM or 25 µM APR-246 (Sigma-Aldrich, SML1789) or 0.01 µM or 0.04 µM ZMC-1 (Abcam, NSC319726, #A24132) alone or in combination with 10 µM N3a or DMSO for a total of 8 days. Starvation experiments were performed to investigate the effect of nutrient deprivation on cell growth. Three different conditions were used to induce starvation: HBSS (Gibco, 14025050) for 3 days, DMEM without glucose (Gibco, 14025050) for 1 day, and DMEM without glutamine (Gibco, 21969) for 7 days. Following starvation, cells were allowed to recover and expand for either 1 day in the case of the -glucose condition or 7 days in the case of the HBSS or -glutamine conditions.

For generating exon-wide mutant cell libraries, 5.4 x 10^8^ HCT116 LSL/ΔIn5 or HCT116 LSL/ΔIn5+7 cells were transfected with 1.125 mg pX459_blast_In5^LSL^ or pX459_blast_In7^LSL^, and 1.125 mg of the corresponding HR700 vector library using Lipofectamine 2000. Transfected cells were selected with 20 µg/ml blasticidin for 3 days and 1 µg/ml puromycin for 7 days. 1.2 x 10^8^ cells were infected with AV-Cre and, after 5 days, the cell library was divided and treated with 10 µM N3a or DMSO for 8 days. Recombination was monitored through FACS-analysis of GFP expression.

### Genomic DNA analysis of single mutants and mutant libraries

Genomic DNA of mutant cells was isolated using DNA Blood Mini Kit (QIAGEN, 51106) following the manufacturer’s protocol, and a nested PCR strategy was used to selectively amplify either edited or edited and recombined alleles (Supplementary Fig. 1a). The input amount of genomic DNA and number of PCR reactions were adjusted to achieve a minimum average coverage of 1000 cells/variant. For first step PCR, the following primers were used before AV-Cre recombination: LoxP fw 5’-GGG GGC TGT CCC TAG ATC TAT AA-3’, Intron 9 rev 5’-GTA TGC CTG TGG TCC TAG CT-3’, and after AV-Cre recombination: Intron 4 fw 5’-CCC TTT GGC TTC CTG TCA GTG, Intron 9 rev 5’-GTA TGC CTG TGG TCC TAG CT-3’. The PCR products were pooled, purified and diluted 1:1000 for the second step. Editing-specific primers were used for the second step PCR, exon-wise, with the following primer pairs: Exon 5 fw 5’-TTG CTT TAT CTG TTC ACT TGT GCC C-3’, Exon 5 rev 5’-CAG TGA GGA ATC AGA GGC CTC C-3’, Exon 6 fw 5’-TGC CCA GGG TCC GGA GGC-3’, Exon 6 rev 5’-GGA GGG CCA CTG ACA ACC ACC C-3’, Exon 7 fw 5’-TCC CCT GCT TGC CAC AGG-3’, Exon 7 rev 5’ -GGA GGA GAA GCC ACA GGT TAA GAG-3’, Exon 8 fw 5’-GCT TTG GGA CCT CTT AAC CTG TG-3’, Exon 8 rev 5’-CAT AAC TGC ACC CTT GGT CTC C-3’. The PCRs were performed with Q5 High-Fidelity DNA Polymerase (New England Biolabs, M0491) following the manufacturer’s protocol. PCR products were purified using PCR purification kit according to the manufacturer’s protocol. PCR amplicons were purified with AMPure XP beads (Beckman Coulter, A63880) and sequencing libraries were prepared from 5 ng of the purified amplicon using the NEBNext Ultra DNA Library Prep Kit for Illumina (New England Biolabs, E7370L) according to manufacturer’s protocol. The quality of sequencing libraries was validated with a Bioanalyzer 2100 using the Agilent High Sensitivity DNA Kit (Agilent, 5067-4626). The pooled sequencing libraries were quantified and sequenced on either the MiSeq (v2 or v2 nano, 2×250 cycles, or v3, 2×300 cycles, depending on library complexity) or NovaSeq (SP 2×250 cycles) platform (Illumina).

Sequenced reads were demultiplexed using an in-house demultiplexer package mmdemultiplex (version 0.1). Overlapping paired-end reads where trimmed of adapter/primer sequences using CutAdapt^62^ (version 3.5) and merged into a single sequence using NGmerge^63^ (version 0.3), taking advantage of the overlapping reads to reduce sequencing errors. The occurrence of each synthetic sequence was counted via exact matching, since the minimal hamming distance between synthetic sequences was 1. To calculate the relative frequencies (variant abundances), the count was divided by the total number of matched reads, excluding any wild-type reads and duplicate sequences. Enrichment scores (ES) were determined as the negative log2 fold change of the variant abundance in treated versus control conditions. However, it should be noted that the ES is dependent on the relative amount of wild-type-like and loss-of-function variants in a cell population, which varies between different libraries. To obtain a score that is comparable across different libraries and screens, the ES was into a relative fitness score (RFS) by scaling the median ES of synonymous variants to -1 and the median ES of nonsense variants to +1 for each library and screen.

### cDNA analysis of single mutants and mutant libraries

Total RNA was isolated using the RNeasy Mini Kit (QIAGEN, 74106) and reverse transcribed using the SuperScript VILO cDNA Synthesis Kit (Invitrogen, 11754250). PCR was performed with the primers: Exon 2 fw 5’-ATG GAG GAG CCG CAG TCA GAT-3’, Exon 11 rev 5’-TCA GTC TGA GTC AGG CCC TTC-3’. PCR products were purified and sequenced using Sanger sequencing. For cDNA sequencing of mutant cell libraries, RNA was reverse transcribed and amplified in 5 reactions with 1 µg RNA template using SuperScript IV One-Step RT-PCR System with ezDNase (Invitrogen, 12595025) and the primers: Intron 4 fw 5’- TCC CAG AAT GCC AGA GGC TGC T-3’, Intron 8 rev 5’-GCT CAC GCC CAC GGA TCT GAA G-3’. The amplified PCR products were pooled, providing an estimated variant coverage of 50-250x, purified and diluted 1:1000 for a second step of PCR, using exon-specific primers and Q5 High-Fidelity DNA Polymerase: Exon 5 fw 5’-TGG GAC AGC CAA GTC TGT GAC T-3’, Exon 5 rev 5’- AGA TGC TGA GGA GGG GCC AGA C-3’, Exon 6 fw 5’-CAT GAC GGA GGT TGT GAG GCG C-3’, Exon 6 rev 5’-TTC ATG CCG CCC ATG CAG GAA C-3’, Exon 7 fw 5’-GTC TGG CCC CTC CTC AGC ATC T-3’, Exon 7 rev 5’ -GGA CAG GCA CAA ACA CGC ACC T-3’, Exon 8 fw 5’-TAA CAG TTC CTG CAT GGG CGG C-3’, Exon 8 rev 5’-GCT GGG GAG AGG AGC TGG TGT T-3’. Library preparation, sequencing, and analysis followed the same protocol as for genomic DNA. Merged reads were trimmed to only include exonic regions.

### RNA-Sequencing

For RNA-Sequencing experiments, cells were treated either with 10 µM N3a or the corresponding volume of DMSO, as solvent control, for 36 h prior to RNA isolation using the RNeasy Mini Kit according to the manufacturer’s protocol. RNA quality was evaluated using the Experion RNA StdSens Analysis Kit (Bio-Rad, 700-7103). RNAseq libraries were prepared from total RNA with the QuantSeq 3′ mRNA-Seq Library Prep Kit FWD for Illumina (Lexogen, 015.24) in combination with the UMI Second Strand Synthesis Module for QuantSeq FWD (Illumina, Read 1) (Lexogen, 081.96) following the manufacturer’s protocol. The Quality of sequencing libraries was validated on a Bioanalyzer 2100 using the Agilent High Sensitivity DNA Kit. Pooled sequencing libraries were quantified and sequenced on the NextSeq 550 platform (Illumina) with 75-base single reads. Unique molecular identifiers (UMI) were extracted and the first four nucleotides corresponding to the QuantSeq FWD-UMI 3’ spacer were removed. The trimmed reads were mapped to the Homo sapiens (revision 104, GRCh38) Ensembl reference genome using STAR^64^ (version 2.7.10a). The UMIs were then deduplicated using UMI-tools^65^ (version 1.1.1). The UMI per gene was quantified and normalized to counts per million (CPM). Genes with CPM counts below 1 in all samples were considered background noise and discarded. Further analysis was restricted to protein-coding and lincRNA genes. Differential gene expression was analyzed via DEseq2 (version 1.34.0)^66^. The obtained p-values were corrected via Benjamini-Hochberg correction. Genes with log2FC ≥ 1 as well as corrected p-values smaller than 0.05 were considered differentially expressed. Principle component analysis was performed on the corrected count matrix after subjecting it to variance-stabilizing transformation via DEseq2. For heatmaps, expression values were z-transformed and genes were clustered using hierarchical average-linkage clustering based on Euclidean distances. Gene set enrichment analysis was performed using gene sets from the Molecular Signatures Database (MSigDB) and GSEA software^67^ (version 4.2.2). Raw sequencing data was deposited at ArrayExpress, accession number E-MTAB-12734.

### Proliferation/IC_50_ Assay

8 x 10^4^ cells were seeded on 96 well plates (Sarstedt, 83.3925) and treated with various doses of N3a after 24 hours. Cells were imaged every 4 hours using the IncuCyte S3 live-cell analysis system (Sartorius). Confluence data was analyzed using the area under the curve (AUC) to calculate IC_50_ values.

### Apoptosis Assay

Cells and media supernatants were collected, pelleted, and resuspended in Annexin V-APC conjugate (MabTag, AnxA100) diluted in Annexin V binding buffer (BD Biosciences, 556454) according to the manufacturer’s protocol. The suspension was incubated in the dark for 20 minutes at RT, washed in Annexin V binding buffer and analyzed by flow cytometry (BD LSR II Flow Cytometer or BD Accuri C6 Plus Cytometer). For sequencing of apoptotic cells, annexin-V positive or negative cells were sorted with a MoFlo Astrios sorter (Beckman Coulter).

### Reverse transcription quantitative PCR (RTqPCR)

Total RNA was isolated using the RNeasy Mini Kit and reverse transcribed using the SuperScript VILO cDNA Synthesis Kit. The resulting cDNA was used for qPCR using ABsolute qPCR Mix SYBR Green (Thermo Fisher, AB1158B) with the following primers: TP53 fw 5’-GGA GCC GCA GTC AGA TCC-3’, TP53 rev 5’-CAA TAT CGT CCG GGG ACA GC-3‘, GAPDH fw 5’-CTA TAA ATT GAG CCC GCA GCC-3’, GAPDH rev 5’-ACC AAA TCC GTT GAC TCC-3‘. The data was analyzed using the ΔΔC_t_ method with GAPDH as reference.

### Western blot analysis

Cells were lysed in NP-40 Lysis Buffer (50 mM Tris-HCl, 150 mM NaCl, 5 mM EDTA, 2% NP-40, pH 8.0) supplemented with cOmplete ULTRA protease inhibitor cocktail (Roche, 4693124001) and sonicated using a Bioruptor (Diagenode) for 5 minutes. 20-40 µg of protein was separated on NuPAGE 4 to 12% Bis-Tris polyacrylamide gels (Invitrogen, WG1402) using MOPS buffer (Invitrogen, NP0001). Following transfer to Immun-Blot PVDF Membrane (BioRad, 1620177), antigens were detected using the antibodies: p53 (Santa Cruz Biotechnology, sc-126; antibody DO-1), p21 (Santa Cruz Biotechnology, sc-6246), β-actin (Abcam, ab6276). Detection was performed with secondary goat anti-mouse IgG Fc HRP antibody (Invitrogen, A16084) and WesternBright Sirius chemiluminescent HRP conjugate (advansta, K-12043). β-actin was detected using goat anti-mouse Alexa-488 conjugate (Invitrogen, A-11029).

### Stability measurements of recombinant p53 DNA-binding domain variants

Cancer mutations were introduced into a stabilized pseudo wild-type variant of the human p53 DNA-binding domain (residues 94-312; M133L/V203A/N239Y/N268D) that we have routinely used as a framework for biophysical and structural studies in the past^45,68^. Sequences of the pET24a-based expression vectors used are given in Supplementary Table 9. The inserts between the NdeI and EcoRI restriction sites encode for a fusion protein containing an N-terminal hexahistidine tag, the lipoyl-binding domain of the dihydrolipoamide acetyltransferase component of pyruvate dehydrogenase complex from *Bacillus stearothermophilus* (Uniprot entry P11961, residues 2-85), followed by a TEV protease cleavage site and human p53 residues 94-312 with mutations of interest. The different p53 DBD variants were expressed in *E. coli* C41 cells and purified by Ni-NTA column, overnight TEV protease cleavage, followed by affinity chromatography on a heparin column and size-exclusion chromatography as described previously^45^.

Melting temperatures, *T*_m_ values, of the purified p53 variants were determined by differential scanning fluorimetry using an Agilent MX3005P real-time qPCR instrument (excitation/emission filters = 492/610 nm). Assay buffer: 25 mM HEPES, pH 7.5, 500 mM NaCl, 0.5 mM TCEP, with a final protein concentration of 5 μΜ and the fluorescent dye SYPRO Orange (Invitrogen, 10338542) at a dilution of 1:1000. The fluorescence signal was monitored upon temperature increase from 25 to 95 °C, at a heating rate of 3 °C/min, and *T*_m_ values were calculated after fitting the fluorescence curves to the Boltzmann function. Measurements were performed in three independent repeats (each consisting of four technical repeats on the same plate). Mutation-induced changes in DBD stability are given as Δ*T*_m_ = *T*_m_ (mutant) – *T*_m_ (wild type) (Supplementary Table 9).

### Animal Models

Mouse experiments were performed in accordance with the German Animal Welfare Law (TierSchG) and received approval from the local authority (Regierungspräsidium Gießen). The mice were housed in specific-pathogen free conditions, kept on a 12-h light/dark cycle and fed a standard housing diet (Altromin, 1328), with access to water ad libitum.

For *in vivo* passaging of HCT116 R175H/Δ cells, 1 x 10^6^ cells were injected intravenously into the tail vein of immunodeficient *Rag2^tm1.1Flv^;Il2rg^tm1.1Flv^* male and female mice that were a minimum of eight weeks old. Starting from the onset of clinical symptoms, mice were monitored daily using a scoring system, and were euthanized by cervical dislocation when critical symptoms appeared or after 12 weeks, whichever came first. Lungs and, if present, metastases were isolated, minced, and incubated in 1 mg/ml Collagenase/Dispase (Roche, 10269638001) and 100 µg/ml DNase I (Roche, 4536282001) at 37 °C for 1 h at 100 rpm on a horizontal shaker. The cells were filtered through a 70 µM EASYstrainer (Greiner Bio-One, 542070), pelleted at 300 x g, and erythrocytes were lysed using red blood cell lysis buffer (150 mM NH_4_Cl, 10 mM NaHCO_3_, 1.27 mM EDTA) for 5 minutes at RT. Cells were washed once with DPBS (Gibco, 14190) and cultured under standard conditions.

For the *in vivo* tumor growth and metastasis assay, HCT116 R175H/Δ p3-met cells were labelled with the intracellular Firefly (FLuc) and secreted Gaussia (GLuc) luciferases using the retroviral plasmid pMSCV_FLuc_T2A_GLuc_Hygro. For plasmid generation, the FLuc ORF was PCR-amplified using the following primers: FLuc_BglII_fw 5‘-AGA TCT CAC CAT GGA AGA TGC CAA AAA CAT TAA G-3‘, FLuc_T2A_rev 5‘-CAC GTC ACC GCA TGT TAG AAG ACT TCC TCT GCC CTC CAC GGC GAT CTT GCC GCC-3‘; the GLuc ORF was PCR-amplified using the following primers: GLuc_T2a_fw 5‘-CTA ACA TGC GGT GAC GTG GAG GAG AAT CCC GGC CCT ATG GGA GTC AAA GTT CTG TTT G-3‘, GLuc_XhoI_rev 5‘-CTC GAG TTA GTC ACC ACC GGC CCC-3‘. Both fragments were fused using overlap extension PCR and cloned into pCR Blunt II-TOPO vector using Zero Blunt TOPO kit (Invitrogen, 450245) according to the manufacturer’s protocol. Lastly, FLuc_T2A_GLuc was cloned into MCS of pMSCV_hygro (Clontech Laboratories, Inc., 631461) using BglII (New England Biolabs, R0144) and XhoI (New England Biolabs, R0146). AmphoPack-293 cells (Takara Bio Inc., CVCL_WI47) were transfected with pMSCV-FLuc-T2A-GLuc-Hygro using calcium phosphate protocol^69^. Three days after transfection, supernatants were collected, filtered using Filtropur S 0.45 (Sarstedt, 83.1826) and supplemented with 8 μg/ml polybrene (Sigma-Aldrich, 83.1826). 2.5 x 10^6^ tumor cells were transduced with retroviral supernatant using spinoculation (1 h, 600 x g, 37°C). To knock-out p53R175H, labelled cells were transfected with pX459_blast with sgTP53_Ex3: 5’- ACT TCC TGA AAA CAA CGT TC-3‘ or sgTP53_Ex5: 5‘-GTT GAT TCC ACA CCC CCG CC-3‘ and selected with 20 µg/ml blasticidin. Growth of primary, subcutaneous tumors was assessed longitudinally by measuring GLuc activity in blood samples as described^70^. Livers were incubated using Luciferase Cell Culture Lysis 5 x Reagent (Promega, E1531) according to the manufacturer’s protocol. Firefly luciferase activity was measured in liver lysates on a plate reader luminometer (ORION II, Titertek-Berthold) using the Beetle-Juice Luciferase assay Firefly (PJK GmbH, 102511-1) according to the manufacturer’s protocol.

### Invasion/Migration assays

1 x 10^5^ cells were seeded in 24-well transwell inserts with 8 µm pore size (Sarstedt, 102511-1) in media containing 1% FBS. The bottom well was filled with medium containing 10% FBS. After 4 days, the cells remaining on the top of the transwell membrane were thoroughly washed using a Q-Tip dipped in DPBS. The transwell inserts were then fixed for 10 min in 70 % EtOH at RT, dried, and stained for 15 min with 0.2% crystal violet (Sigma-Aldrich, HT90132) in 10 % EtOH. The transwell inserts were washed in water and allowed to dry. For quantification, 200 µl of 20% acetic acid was added to each well and the dish was shaken for 15 min at 350 rpm at RT. The solution was then collected into 96-well plates and the absorbance was measured at 590 nm in a CYTATION 3 imaging plate reader. For normalization, 24-wells were seeded with the same number of cells in media with 1 % FBS and stained one day after plating.

For invasion assays, the transwell inserts were pre-coated with 50 µl Matrigel (Corning, 354234) prior to seeding the cells. Before staining, the Matrigel was removed using a Q-Tip, and the upper part of membrane was thoroughly washed with DPBS.

In the case of siRNA transfections, the cells transfected with the ON-TARGETplus siRNA set of 4 (Horizon Discovery Ltd., LQ-003329-00 for *TP53* or D-001810-10 as non-targeting control) using Lipofectamine RNAiMax (Thermo Fisher Scientific, 3778075) according to the manufacturer’s protocol and plated on the transwell inserts 24 hours after siRNA transfection.

### Data analysis and Software

To evaluate the correlation between RFS value and variant frequency in cancer patient samples, we obtained *TP53* mutation data from the following public databases: UMD TP53 Mutation Database^2^ (release 2017_R2, https://p53.fr/tp53-database), NCI/IARC The TP53 Database^71^ (release R20, July 2019, https://tp53.isb-cgc.org/), the ‘curated set of non-redundant studies’ from the TCGA and the AACR project GENIE^72^ from cBioPortal^73^ (http://www.cbioportal.org/, downloaded on Dec 20, 2022). To evaluate the mutational probability of variants, we used Mutational Signatures (v3.3 – June 2022) downloaded from COSMIC (https://cancer.sanger.ac.uk/signatures/)34. We calculated a mean Single Base Substitution (SBSmean) signature by averaging signatures SBS1 to SBS21weighted by their prevalence in cancer samples, as reported by Alexandrov et al., 2013^74^ (Supplementary Table 3).

The evolutionary conservation profile for p53 was downloaded from the ConSurf-Database (https://consurf.tau.ac.il/consurf_index.php)75. RFS values were mapped onto the p53DBD structure using PyMOL (version 2.5.2) with Protein Data Bank (PDB) entries 2AHI^76^ (https://www.rcsb.org/structure/2AHI) and 3KZ8^77^ (https://www.rcsb.org/structure/3KZ8). To analyze distance relationships within the p53DBD, we generated a contact map for PDB 2AHI using ProteinTools (https://proteintools.uni-bayreuth.de/)78. The map represents the distances between all amino acid pairs in a matrix form. The distance from the DNA-binding surface (TOP) was defined as the mean distance from the residues 248, 273, 277, and 280; the distance from the opposite pole (BOTTOM) as the mean distance from residues 153, 225 and 260; and the distance from the core (CENTER) as the mean distance from residues 195, 236, and 253. HoTMuSiC^39^ was used to predict thermal destabilization of variants and solvent accessibility of residues based on PDB entry 2AHI.

To compare CRISPR and cDNA-based mutome screens, we used datasets from Kotler et al. 2018^8^ and Giacomelli et al., 2018^4^. From Kotler et al., we used enrichment data of p53 variants measured in the p53-null H1299 cell line (Supplementary Table 2, RFS_H1299), and from Giacomelli et al., we used enrichment results from the A549 p53-knockout cell line (A549_p53NULL_Nutlin-3_Z-score). Both cDNA datasets were transformed to RFS as defined above by scaling the median of nonsense variants to +1 and the median of synonymous variants to -1. We used yeast reporter data from Kato et al., 2003^7^ (downloaded from the NCI/IARC TP53 Database) to analyze transcriptional activity of p53 variants. Our analysis used the mean transcriptional activity (in % of wild type) from 8 different reporter constructs.

The plots and statistical analyses in this study were created using GraphPad Prism (version 9.4.1) or Python (version 3.9.12) with libraries: Matplotlib (version 3.5.1), Seaborn (version 0.11.2), SciPy (version 1.7.3), and Statsmodels (version 0.13.2). Graphics were assembled in Adobe Illustrator (version 26.5.2). The results presented in the graphs represent the mean or median values obtained from n biological replicates. The error bars in the figures indicate the standard deviation (SD), unless stated otherwise. The difference between two sets of data was assessed through either a two-sided unpaired t-test or a Mann-Whitney test if the data was not normally distributed. To analyze multiple groups, a one-way ANOVA was used in combination with a multiple comparisons test. For three or more groups that have been divided into two independent variables (such as treatment and genotype), a two-way ANOVA was used in combination with a multiple comparisons test. The ANOVA results and selected pairwise comparisons are reported in the figures and source data files. A p-value less than 0.05 was considered statistically significant.

## Data availability

All data generated or analyzed during this study are included in this published article (and its supplementary information files). RNA sequencing data was deposited at ArrayExpress, accession number E-MTAB-12734.

## Oligonucleotides

All oligonucleotide sequences are provided in Supplementary Table 11.

## Acknowledgements

The authors thank Sigrid Bischofsberger, Antje Grzeschiczek, Björn Geißert, Angelika Filmer, Alexandra Schneider and employees of the animal facility of Marburg University for technical and experimental support. We acknowledge support by the Medical Core Facilities for Genomics and Flow Cytometry. D.-I.B. and A.C.J. are grateful for support by the Structural Genomics Consortium (SGC), a registered charity (No:1097737) that received funds from Bayer AG, Boehringer Ingelheim, Bristol Myers Squibb, Genentech, Genome Canada through Ontario Genomics Institute, Janssen, Merck KGaA, Pfizer, and Takeda. A.C.J. is funded by German Research Foundation (DFG) grant JO 1473/1-3. T.S. is funded by grants from BMBF (031L0063 to TS), Deutsche Forschungsgemeinschaft (TRR81/3 109546710 Project A10, STI 182/13-1, STI 182/15-1, GRK2573), German Center for Lung Research (DZL), State of Hesse (LOEWE iCANx), von Behring-Röntgen Stiftung (65-0004 and 66-LV06), Deutsche José Carreras Leukämie Stiftung e.V. (09 R/2018). The funding bodies were not involved in the design of the study and collection, analysis, and interpretation of data and in writing the manuscript.

## Author contributions

The study was conceptualized by T.S. with support from J.F., M.K., R.S., and A.C.J. The wet-lab experiments were performed by J.F., M.K., E.P., M.N., D.D., M.N., P.H., A.B., D.-I.B., K.K., N.M., and I.B. The animal experiments were conducted by J.F., M.K., E.P., and S.E. The experiments were supervised by M.W., S.E. A.C.J., and T.S. Next generation sequencing was performed by A.N., T.P., and M.B. The data was curated, analyzed, and visualized by K.H., M.M., J.F., M.K., and T.S. Funding was acquired by R.S., A.C.J., and T.S. The original draft was written by J.F. and T.S., and all co-authors reviewed and edited the manuscript.

## Supplementary Figure Legends

**Supplementary Fig. 1. Generation and functional characterization of single *TP53*-mutant HCT116 cell clones. a**, CRISPR/Cas9-mediated targeting of *TP53* in HCT116 cells. Shown are the two *TP53* alleles in HCT116 cells and their modifications in HCT116 LSL/Δ cells. Exon 5 and 6 mutations were engineered in the cell line depicted on the left. For mutagenesis of exon 7 and 8, this cell line was further modified as shown on the right. The Δ allele contains inactivating deletions in introns 5 and 7 as indicated. The second allele contains a loxP-flanked transcriptional stop (LSL) cassette expressing GFP and a non-functional Puromycin N-acetyltransferase (Puro^mut^) resistance gene in intron 4. In addition, the LSL allele harbors alterations (SNV in intron 5, deletion in intron 7) to enable specific Cas9-targeting of the LSL allele only. Donor vectors, used to introduce mutations, contain an intact Puromycin resistance gene allowing selection of HDR-edited cells by Puromycin treatment. To prevent re-cutting and selective PCR amplification of HDR-edited alleles (LSL-mut), donors for targeting of exons 5/6 contain an additional PAM-inactivating mutation. In case of targeting exons 7/8, a deletion in intron 7 on the LSL allele eliminated the need for an additional donor mutation. Adenoviral Cre infection was used to excise the LSL-cassette and activate expression of the mutated allele, yielding HCT116 mut/Δ cells. Selective next-generation sequencing of the edited and Cre-recombined allele was ensured by a nested PCR amplification strategy using the indicated, color-coded, primer pairs. **b** and **c**, Sanger sequencing of intron 5 and 7 deletions in HCT116 LSL/Δ cells. **d**, Western blot validating Cre and N3a-dependent mutant p53 expression, using LSL-WT and LSL-R175H cells as an example. **e**, Proliferation of additional *TP53*-mutant cell clones in presence of increasing concentrations of N3a analyzed by real-time live cell imaging. Shown is the area under the proliferation curve (AUC) relative to untreated. **f**, Western blot demonstrating low variability in mutant p53 expression between different R175H cell clones. **g-h**, Proliferation of different R175H cell clones in presence of increasing concentrations of N3a analyzed by real-time live cell imaging. p53-null (LSL, red) and wild-type (WT, green) are shown for reference. **g**, Area under the proliferation curve (AUC) relative to untreated. **h**, 50% inhibitory concentration (mean IC_50_ and 95% CI, n=3).

**Supplementary Fig. 2. Impact of R175 variants on drug responses. a**, Dose-dependent changes in R175 variant abundance following 8 days of N3a treatment. Heatmap shows enrichment (or depletion) as the -log2 fold change versus the mean of the DMSO-treated control. n=3 biological replicates per condition. **b-c**, Impact of R175 variants on the cellular response to 8-day treatment with the indicated concentrations of APR-246 and N3a. **b**, Heatmap depicting changes in variant abundance as -log2 fold change versus the untreated control. n=3 biological replicates per condition. **c**, Heatmap showing pair-wise correlation coefficients (ρ, Spearman). Dendrogram shows hierarchical clustering of samples using average linkage and Euclidean distance. **d-e**, Impact of R175 variants on the cellular response to 8-day treatment with the indicated concentrations of ZMC1 and N3a. **d**, Heatmap depicting changes in variant abundance as -log2 fold change versus the untreated control. n=3 biological replicates per condition. **e**, Heatmap showing pair-wise correlation coefficients (ρ, Spearman). Dendrogram shows hierarchical clustering of samples using average linkage and Euclidean distance.

**Supplementary Fig. 3. R175 mutagenesis screen in H460 cells. a**, Scheme depicting generation of the H460 LSL/Δ/Δ cell line. **b**, Sanger sequencing results of the three *TP53* alleles in H460 LSL/Δ/Δ cells. **c**, Sanger sequencing results of the LSL allele at codon 175 for R175-edited/mutated H460 cell clones. **d**, Western blot of mutated H460 cell clones ± Cre and N3a. **e**, Heatmap depicting changes in variant abundance following 8 days of 10 µM N3a treatment. Shown is the -log2 fold change versus the untreated control for n=3 (HCT116) and n=6 (H460) biological replicates. **f**, Scatter plot illustrating the correlation between N3a-induced variant enrichment in HCT116 and H460 cells. Shown is the mean ±SD enrichment (-log2 FC, n=3 biological replicates) and Pearson correlation coefficient ρ. Dashed line, line of identity.

**Supplementary Fig. 4. Codon 176-185 mutagenesis screen. a-c**, Quality control plots illustrating correlation of variant abundance between donor (plasmid) library and variant cell libraries before and after Cre recombination (-Cre and +Cre) and following 8 days of 10 µM N3a treatment (+N3a). Shown is the mean abundance (n=3 biological replicates) for synonymous (syn, green), null (red) and missense (mis, grey) variants. Dashed line, line of identity. **d**, Heatmap showing pair-wise correlation coefficients (ρ, Spearman). Dendrogram shows hierarchical clustering of samples using average linkage and Euclidean distance. **e**, Heatmap of variant enrichment/depletion after 8 days of treatment with indicated concentrations of Mdm2/Mdmx inhibitors. Shown is the -log2 FC relative to the mean of DMSO-treated control cells for n=3 biological replicates. **f**, Heatmap of variant enrichment after 8 days of 10 µM N3a treatment. Shown is the mean -log2 FC relative to DMSO-treated control cells. Synonymous mutations are marked with a yellow dot.

**Supplementary Fig. 5. *TP53* DBD mutome screen. a**, Quality control plots illustrating correlation of variant abundance between donor (plasmid) library and variant cell libraries before and after Cre recombination (-Cre and +Cre) and following 8 days of 10 µM N3a treatment (+N3a). Shown is the median abundance (n=3 biological replicates) for synonymous (syn, green), nonsense (non, red) and other (grey) variants. ρ, Spearman correlation coefficient. Dashed line, line of identity. **b**, Kernel density estimation (KDE) plots of variant abundance in indicated donor and cell libraries. **c**, KDE plot of the log2 fold change (FC) of variant abundance under N3a treatment. **d**, KDE plot of the relative fitness score (RFS) of variants. **e**, Bar plot showing the median RFS values of all perturbations at exemplary codons (blue, negative RFS indicative of WTp53-like activity; red, positive RFS indicative of loss of WTp53 function). Black bars indicate the number of patients with the respected amino acid substitution in the UMD *TP53* mutation database. **f**, Hierarchically clustered heatmap showing the RFS for all mis, syn, and non variants. Exemplary codons (mutational hotspots and codons from **e**) are labelled. Bar plots show for each codon the mutation frequency in the UMD *TP53* mutation database, the evolutionary conservation score, and the RFS (mean±SD) of all missense substitutions at this position. **g**, Scatter plots showing the correlation between RFS and distance of the altered residue from the TOP (DNA-binding surface), BOTTOM (protein pole opposite from the DNA-binding surface) and CENTER of the p53 DBD. **h-j**, Scatter plots showing the correlation between RFS and (**h**) solvent accessibility of the altered residue, (**i**) thermal destabilization of the variant as predicted by HoTMuSiC^39^, and (**j**) the conservation score of the altered residue (ConSurf Database^75^). In **h**, solvent-accessible residues with nevertheless high RFS values are indicated. R248 is a DNA contact residue, E224 and S261 are located at exon borders and affect splicing, G199 is located at the inter-dimer interface and also critical for splicing. **g-j**, All plots show variants as individual datapoints, kernel density estimates, and regression lines with 95% confidence intervals. ρ, Spearman correlation coefficient.

**Supplementary Fig. 6. RFS and mutational probability. a-d**, Scatter plots of RFS versus patient count (sum of all records in the UMD, IARC/NCI, TCGA, and GENIE databases). **a**, Variants are colored by mutation class. Labelled in red are functionally neutral genetic variants (polymorphisms, poly) according to Doffe et al., 2020^32^. **b**, Missense variants colored by the number of substituted nucleotides. **c**, Single-nucleotide missense variants colored as transition Ts (A-G, C-T) or transversion Tv (A-C, A-T, C-A, C-G) mutations. **d**, Single-nucleotide missense variants colored as CpG or non-CpG mutations. **e-h**, Violin plots showing the distribution of patient counts for the mutation types depicted in **a-d** stratified by RFS as RFS+ (RFS>0) or RFS- (RFS<0). n.o., not observed. Tables report the two-way ANOVA p-value and effect size (ω^2^) for each factor and their interaction. Selected post-hoc multiple comparison test results (Tukey) are shown directly in the plot. **i-j**, Scatter plots of RFS versus patient count (sum of all records in the UMD, IARC/NCI, TCGA, and GENIE databases) colored by mutational probability according to the indicated COSMIC mutational signatures (v3.3 - June 2022)^34^. SBSmean (**j**) denotes an averaged mutational signature calculated by weighting the most common mutational signatures based on their occurrence in the TCGA pan-cancer cohort^74^. **k** and **l**, Violin plots comparing the distribution of patient counts for single-nucleotide substitutions stratified by RFS as RFS+ (RFS>0, left violin half) or RFS- (RFS<0, right violin half). **k**, All single-nucleotide substitutions and p-value from a two-sided Mann-Whitney test. Lines show the median and the 25% and 50% quartiles. **l**, Single-nucleotide substitutions binned by increasing mutational probability using the ‘SBSmean’ signature. Two-way ANOVA p-value and effect size (ω^2^) for each factor (‘RFS’ and ‘SBSmean bin’) and their interaction are reported in the table, indicating a strong effect of RFS on patient count mostly independent of mutational probability.

**Supplementary Fig. 7. Selection of CRISPR-identified LOF variants during tumorigenesis. a**, Scatter plots illustrating correlation between RFS values obtained by CRISPR mutagenesis and cDNA overexpression (Kotler et al., 2018^8^). Variants are colored based on their mutational probability according to the indicated COSMIC mutational signatures (v3.3 - June 2022)^34^. **b**, Violin plots depict the distribution of mutational probabilities among the variants located in the three main quadrants LL, LR and UR. Reported are p-values from Tukey’s post-hoc multiple comparisons tests performed after one-way ANOVA. **c**, Scatter plots illustrate the correlation between RFS values obtained by CRISPR mutagenesis and cDNA overexpression (Kotler et al., 2018^8^). Variants are colored by their frequency in patients based on the indicated mutation databases (UMD, IARC/NCI, TCGA, GENIE). Violin plots depict the distribution of variant patient counts in the three main quadrants LL, LR and UR (p-values from one-way ANOVA and Tukey’s post-hoc multiple comparisons tests). All violin plots show the median and the 25% and 50% quartiles.

**Supplementary Fig. 8. Comparison of CRISPR and cDNA overexpression screening results. a-f**, Scatter plots illustrate the correlation between RFS values obtained by CRISPR mutagenesis and cDNA overexpression (Giacomelli et al., 2018^4^). Variants are colored based on (**a**) mutation type, (**b**) thermal stability as predicted by HoTMuSiC^39^, (**c**) average mutational probability using a weighted average mutational signature SBSmean, (**d**) frequency in cancer patients (sum of all records in UMD, IARC/NCI, TCGA, GENIE databases), (**e**) mean transcriptional activity relative to wild-type p53 (WT) as measured in a yeast-based reporter system (Kato et al., 2003^7^), and (**f**) classification as pLOF (20-60% transcriptional activity). Inserted violin plots illustrate the value distribution in the four quadrants (LL, lower left; LR, lower right; UL, upper left; UR, upper right) and report p values from one-way ANOVA with Tukey’s post-hoc multiple comparisons tests (***, <0.001; ns, not significant).

**Supplementary Fig. 9. Missense and synonymous mutations resulting in splicing defects. a-d**, Scatter plots comparing the abundance of missense variants in the cell libraries at the level of genomic DNA and mRNA. Each dot represents the median abundance of a variant from n=3 biological replicates. Variants are colored by RFS in **a**, by distance from the exon border in **b**, substitution type in **c**, and patient count in **d**. Variants that are recurrently observed in cancer patients showing evidence for NMD (i.e., underrepresented at the mRNA level) are individually labeled (black font for variants at the exon border, red font for variants inside the exon). Dashed line, line of identity. **e-g**, Schematic depiction of splicing alterations caused by (**e**) L137Q, (**f**) G199G (GGA>GGT) and (**g**) g.7673847 T>G. **h**, Abundance of the regularly spliced and alternatively spliced mRNAs compared to the abundance of the indicated variants at the level of genomic DNA. Alternatively spliced mRNA with in-frame deletions/insertions caused by L137Q and g.7673847 T>G are well detectable and more abundant than the regularly spliced transcript. The alternatively spliced (frameshifted) mRNA resulting from G199G (GGA>GGT) is detectable, but underrepresented compared to its genomic abundance, indicating its clearance by NMD.

**Supplementary Fig. 10. R175H enables evolutionary development of pro-metastatic properties. a**, Images of transwell migration and invasion assays of indicated HCT116 R175H/Δ cells. The top three rows show migrated/invaded cells after crystal violet staining. The bottom three rows show wells with equal amounts of seeded cells as proliferation controls. **b,** Proliferation curves of R175H p0 and p3-met cells measured by real-time live cell imaging. Shown is the mean confluence of n=3 experiments. **c**, Images of transwell migration and invasion assays of indicated HCT116 R175H/Δ and HCT116 R175X/Δ cells transfected with non-silencing (nsi) or p53-depleting siRNA (p53si). The top three rows show 3 replicate wells of migrated/invaded cells after crystal violet stain. The bottom three rows show 3 replicate wells with equal amounts of seeded cells as proliferation controls. **d-h**, Generation and validation of HCT116 R175H/Δ p3-met cells with CRISPR-knockout of p53^R175H^. **d**, Proliferation of HCT116 R175H/Δ cells after transduction with p53-targeting or control Cas9-nucleases. **e**, Western blot confirming efficient p53-knockout. **f**, Images of three replicate wells of migrated and invaded cells after crystal violet staining. **g** and **h**, Quantification of migration (**g**) and invasion (**h**). Shown is the mean ±SD (n=3 replicates) relative to sgCtrl-cells and results from one-way ANOVA with Dunnett’s multiple comparisons test. **i-k**, Transwell migration assays of indicated HCT116 R175H/Δ and HCT116 R175X/Δ p0 cells treated with Nutlin-3a (N3a) or DMSO (D) as control. **i**, Western blot. **j**, Images of three replicate wells of migrated cells stained with crystal violet. **k**, Quantification of migration. Shown is the mean ±SD (n=3 replicates) relative to DMSO-treated R175X cells and results from one-way ANOVA with Dunnett’s multiple comparisons test.

